# Neuropathic pain drives time-dependent reorganization of naturalistic movement and corticostriatal circuits

**DOI:** 10.1101/2025.10.07.680927

**Authors:** Arlene J. George, Ivan Linares-Garcia, Alex J. Yonk, Xinyi C. Zhang, Justin Burdge, Ammar Danish, Mark Gradwell, Victoria E. Abraira, David J. Margolis

## Abstract

Chronic pain fundamentally alters sensorimotor behavior and naturalistic movement, leading to debilitating changes in quality of life. While human neuroimaging studies suggest that chronic pain leads to functional changes in the dorsal striatum (DStr), a region imperative for shaping action selection and movement patterns, most preclinical studies have focused on ventral striatum’s role in reward and motivation, leaving critical questions about DStr circuitry and chronic pain-related behavior unresolved. Furthermore, the reliance on standard evoked reflexive assays misses the persistent, complex behavioral adaptations most relevant to patient wellbeing. Here, we combine machine learning for detailed, automated behavioral characterization with *in vivo* fiber photometry and *ex vivo* synaptic physiology in the spared nerve injury (SNI) mouse model of chronic pain. Through this longitudinal approach, we identified specific functional changes in DStr circuitry that define distinct temporal stages of chronic pain progression. At 3 weeks post-injury, a critical “transition phase”, patch clamp recordings revealed transiently depressing synaptic inputs from primary sensory cortex (S1) to D2-expressing spiny projection neurons (SPNs). Concurrently, unsupervised machine learning demonstrated that behavioral syntax, the structure and sequencing of movement, diverged most sharply during this 3-week window. Conversely, by the late chronic phase at 12 weeks, longitudinal photometry revealed a profound reorganization of D1-SPNs around naturalistic behaviors like grooming and rearing. These findings uncover a pathway-specific corticostriatal mechanism for pain chronification, identifying the DStr as a key mediator of pain-induced movement reorganization and highlighting the 3-week transition phase as a critical therapeutic window to intervene in the progression of chronic pain.

## Introduction

Chronic pain is associated with persistent changes in both sensory processing and motor behavior. Whereas acute pain can produce adaptive changes in movement that protect injured tissue, persistent pain is associated with longer-lasting alterations in movement strategies, locomotor activity, and action selection ^1–5^. Chronic pain is a multidimensional nervous system disease driven collectively by sensory, psychological, and social factors, along with also being a sensorimotor phenomenon, a disorder of how movement is selected and executed. The dorsal striatum (DStr), a core component of the basal ganglia circuit, is the convergence point for sensory and motor signals to select, initiate, and refine motor actions, making it a critical candidate for mediating pain-induced behavioral adaptations. This subcortical structure also receives direct input from the primary somatosensory cortex (S1)^6^, a cortical terminus of the spinothalamic tract that carries pain information^7^. Notably, chronic pain conditions exhibit hyperactivity in the S1 region^8–10^, pointing to the critical role of S1 inputs in driving pain-related behaviors and suggesting vulnerability of the S1-DStr pathway to this sustained hyperactivity. Clinical neuroimaging has shown that the human brain undergoes extensive functional and morphological reorganization of many different brain areas including cortical, limbic, and thalamic regions, that develop during persistent pain^11, 12^. In fact, human neuroimaging studies have consistently identified the corticostriatal circuitry, a key area of movement and action selection function, as the site where pain transitions from acute to chronic stages^13–15^. Furthermore, patients with DStr diseases especially those with motor dysfunction symptoms such as Parkinson’s and Huntington’s disease, also have sensory deficits and pain processing abnormalities^16–18^, pointing to the role of the striatum in pain modulation. Chronic pain produces altered sensory processing and changes in movement, positioning the DStr circuitry as a compelling but understudied network in chronic pain pathology.

Clinical evidence suggests not only is the striatum active during sustained pain, but that pain modulation is mediated by dopamine and specific striatal cell types, namely dopamine D2 receptors^19, 20^. The majority of the striatum consists of spiny projection neurons (SPNs) (∼95%), dopamine receptor 1- and 2-expressing SPNs (D1-and D2-SPNs), which mediate the direct and indirect pathways, respectively. There is a body of striatal research that has defined the complex roles of D1- and D2-SPNs in controlling behavior and movement that range from D1- and D2-SPNs having opposing actions^21, 22^ to being coactive during behavior^23, 24^. Recent evidence has emerged establishing the role of the D2 pathway in the suppression of certain behaviors^25^. In relation to pain, preclinical studies have also linked D2 population function within the DStr to pain behavior, but only in acute nociceptive models^26–28^. This functional organization suggests that chronic pain might differentially affect both D1 and D2 populations, potentially shifting the balance toward avoidance behaviors that characterize chronic pain states. However, how this circuitry drives the behavioral transition from acute to chronic pain remains unknown.

When the pain experience persists for longer than 3 months, chronic pain becomes maladaptive, leading to reorganized movement strategies that perpetuate the chronic pain condition, hinder recovery, and diminish quality of life^29–32^. The experience of pain is inherently multifactorial, shaped by sensory, psychological, and motor factors. However, both clinical and preclinical pain researchers often study these factors in isolation or in limited combinations, leaving a large gap in knowledge of how they collectively influence pain processing and recovery^33^. Chronic pain intrinsically alters how we move and interact with our environment, but much of the preclinical pain research has extensively defined hypersensitivity almost exclusively through standard, evoked reflexive assays and limited to only 2-3 weeks post-injury. By focusing purely on reflexive responses to painful stimuli, these studies overlook the broader adaptation and feedback of overall movement that occur during pain processing at longer timescales. The clinical literature agrees that chronic pain shifts naturalistic movement including gait, posture, and general locomotor activity long after the tissue has healed^34, 35^. Therefore, integrating these complex sensorimotor adaptations is critical for informing evidence-based treatments.

Here, we hypothesized that the transition from acute to chronic pain produces distinct, time-dependent changes in D1- and D2-SPN population activity and drives plasticity in S1-DStr circuits. Using the spared nerve injury (SNI) model, we tracked striatal circuit function across acute (week 1), transition (week 3), and late chronic (weeks 12) phases of neuropathic pain. To our knowledge, this is the first longitudinal study to investigate the DStr in a preclinical neuropathic pain model using a multidimensional approach that integrates both automated evoked reflex kinematics and naturalistic behavior tracking. Through this longitudinal approach, we identify specific functional changes in DStr circuitry that temporally align with critical stages of chronic pain progression. Utilizing an automated reproducible mechanostimulator (ARM), we first established that persistent mechanical hypersensitivity becomes uncoupled from withdrawal vigor across the chronic timescale. During 12-week longitudinal photometry measurements, we found differential engagement of striatal output pathways, culminating in substantial D1 signaling reorganization around ethologically relevant behaviors such as grooming and rearing. *Ex vivo* patch clamp recordings revealed transiently depressing synaptic inputs of S1 to D2 SPNs at 3 weeks post-SNI. Concurrently, unsupervised machine learning of naturalistic behaviors demonstrated that the behavioral syntax, the structure and sequencing of movement, diverged most sharply at Week 3, mirroring this synaptic instability. Collectively, this multidimensional approach allows us to functionally define Week 3 as a critical “transition phase” bridging acute and late chronic pain. By moving beyond traditional evoked assays and utilizing automated machine learning to classify complex behaviors, this study identifies both novel functional roles for the DStr and a critical early-chronic window (Week 3) that could be a critical time point for therapeutic interventions before the maladaptive transition to chronic pain.

## Methods and Materials

### Experimental Animals and Genetic Models

All experimental procedures involving animals including housing, surgical interventions, behavioral assays, and euthanasia were reviewed and approved by the Rutgers University Institutional Animal Care and Use Committee (IACUC Protocol #: 999900197) in accordance with the NIH Guide for the Care and Use of Laboratory Animals. Mice were housed in a temperature- and humidity-controlled facility on a 12:12 h reverse light/dark cycle (lights off at 07:00) with food and water available ad libitum. All handling and behavioral assays were conducted during the active dark phase. To genetically target and differentiate striatal direct-pathway and indirect-pathway spiny projection neurons (D1-SPNs and D2-SPNs), we utilized heterozygous BAC transgenic mouse lines: D1-Cre: B6.FVB(Cg)-Tg(DrD1-cre)EY262Gsat/Mmucd (MMRRC, Cat# 030989) and A2a-Cre: B6.FVB(Cg)-Tg(Adora2a-cre)KG139Gsat/Mmucd (MMRRC, Cat# 036158). For histological validation and targeted patch-clamp electrophysiology, Cre driver lines were crossed with the Cre-dependent fluorescent reporter line Ai14 (B6.Cg-Gt(ROSA)26Sor^tm^^14(CAG-tdTomato)Hze^/J; The Jackson Laboratory, Cat# 007914) to express tdTomato in D1- or A2A-expressing populations. Experimental cohorts comprised male and female mice starting at 8 weeks of age (maintained across experimental timelines up to 12 weeks post-injury).

### Stereotaxic Surgeries and Viral Delivery

Surgeries were executed under aseptic conditions using a digital stereotaxic frame (Stoelting or David Kopf Instruments). Anesthesia was induced at 5% isoflurane in oxygen (1 L/min) and maintained at 1.5–2.0%. Core body temperature was regulated at 35–37°C using a closed-loop homeothermic heating pad (FHC). Ophthalmic ointment (Akorn) was applied to protect the corneas. Pre-operative analgesia was delivered subcutaneously via extended-release buprenorphine (Ethiqa XR, 3.25 mg/kg; Fidelis Animal Health) into the flank. The surgical site was shaved, infiltrated with local bupivacaine (0.03 mg/kg, 0.25%, Fresenius Kabi), and sterilized with three alternating scrubs of Betadine (Purdue Products) and 70% ethanol.

#### Fiber Photometry Cannula Implantation (DLS)

A midline incision was made on the scalp, and a circular piece was removed to expose the skull. The lateral muscles were separated from the skull and the skull was cleaned by scraping away the periosteum. Dental etch bonding agent (iBond; Heraeus Kulzer) was applied to the clean skull and cured with blue light 2x for 20 seconds. Dental composite (Charisma; Heraeus Kulzer) was applied to the outer edge of the skull and cured with blue light 2x for 20 seconds. After ensuring that bregma and lambda coordinates were equal in the dorsoventral plane, one craniotomy was made at the dorsolateral striatum (DLS) (AP = +0.50 mm, ML = -2.50 mm, DV = -2.20 mm). Unilateral AAV injection into the DLS of the right hemisphere was performed first with a micropipette containing GCaMP8s (pGP-AAV-syn-FLEX-jGCaMP8s-WPRE; Addgene #162377-AAV9)^36^ and slowly lowered to the appropriate depth and permitted to sit for 5 min. About 300nL of the GCaMP was automatically injected over the course of ∼15 minutes via the Nanoject III system (Drummond Scientific). After an additional delay of 5 min, the micropipette was slowly raised. A custom fiber optic cannula (FP400ERT, 400um, 0.50 NA mm, stainless steel, Ø2.5 mm ferrule, flat cleave, 2.5mm length, Thor Labs) lowered to the appropriate DV coordinate and secured with dental composite (Tetric Evoflow; Heraeus Kulzer) that was cured with blue light 1x for 20 seconds. The HHMI headpost^37^ was fitted for each mouse and previously described (Yonk et al., 2025) and delicately secured on the posterior area of the charisma ring with cyanoacrylate and dental composite. Finally, a single layer of dental composite was used to build and secure the rest of the headcap before being cured 4x with blue light for 20 seconds each. The scalp was closed around the headcap using cyanoacrylate. Immediately after surgery, mice were placed in a sterile, clean cage that was half-on/half-off a heating pad until ambulation was observed. Mice were monitored for post-operative care for 72 hr. Three weeks post-surgery, mice were handled and habituated to behavioral equipment which also allowed for proper viral expression in the DLS.

#### Corticostriatal Optogenetic Targeting (S1HL to DLS)

A midline incision was made on the scalp to expose the skull and the fascia was cleared away. After ensuring that bregma and lambda coordinates were equal in the dorsoventral plane, one craniotomy was made at the primary somatosensory cortex, hindlimb region (S1HL) (AP = -0.50 mm, ML = -1.40 mm, DV = -0.60 mm). Unilateral AAV injection into S1HL of the right hemisphere was performed first with a micropipette containing Channelrhodopsin-2 (ChR2; pAAV-hSyn-hChR2(H134R)-EYFP; Addgene #26973)^38^ and slowly lowered to the appropriate depth and permitted to sit for 5 min. About 300nL of the ChR2 was automatically injected over the course of ∼15 minutes via the Nanoject III system (Drummond Scientific). After an additional delay of 5 min, the micropipette was slowly raised. The scalp was sutured and secured with tissue glue (Vetbond). Immediately after surgery, mice were placed in a sterile, clean cage that was half-on/half-off a heating pad until ambulation was observed. Mice were monitored for post-operative care for 72 hr. Three weeks post-surgery, mice were handled and habituated to behavioral equipment which also allowed for proper viral expression in the S1HL axon terminals in the DLS.

#### Anterograde tracing (S1HL)

A midline incision was made on the scalp to expose the skull and the fascia was cleared away. After ensuring that bregma and lambda coordinates were equal in the dorsoventral plane, one craniotomy was made at the primary somatosensory cortex, hindlimb region (S1HL) (AP = -0.50 mm, ML = -1.40 mm, DV = -0.60 mm). Unilateral AAV injection into S1HL of the right hemisphere was performed first with a micropipette containing the anterograde tracer (pAAV-CAG-tdTomato (codon diversified); Addgene # 59462) and slowly lowered to the appropriate depth and permitted to sit for 5 min. About 300nL of the virus was automatically injected over the course of ∼15 minutes via the Nanoject III system (Drummond Scientific). After an additional delay of 5 min, the micropipette was slowly raised. The scalp was sutured and secured with tissue glue (Vetbond). Immediately after surgery, mice were placed in a sterile, clean cage that was half-on/half-off a heating pad until ambulation was observed. Mice were monitored for post-operative care for 72 hr. Three weeks post-surgery, mice were handled and habituated to behavioral equipment which also allowed for proper viral expression in the S1HL projections.

#### Retrograde tracing (DLS)

A midline incision was made on the scalp to expose the skull and the fascia was cleared away. After ensuring that bregma and lambda coordinates were equal in the dorsoventral plane, one craniotomy was made at the (DLS) (AP = +0.50 mm, ML = -2.50 mm, DV = -2.20 mm). Unilateral AAV injection into DLS of the right hemisphere was performed first with a micropipette containing the retrograde tracer (pAAV-CAG-GFP; Addgene # 37825) and slowly lowered to the appropriate depth and permitted to sit for 5 min. About 300nL of the virus was automatically injected over the course of ∼15 minutes via the Nanoject III system (Drummond Scientific). After an additional delay of 5 min, the micropipette was slowly raised. The scalp was sutured and secured with tissue glue (Vetbond). Immediately after surgery, mice were placed in a sterile, clean cage that was half-on/half-off a heating pad until ambulation was observed. Mice were monitored for post-operative care for 72 hr. Three weeks post-surgery, mice were handled and habituated to behavioral equipment which also allowed for proper viral expression of inputs to the DLS.

### Spared nerve injury (SNI)

Peripheral neuropathic pain was modeled using the spared nerve injury (SNI) paradigm of the left sciatic nerve branches^39^. Under 1.5–2% isoflurane anesthesia, the lateral surface of the left thigh was shaved and sterilized with three alternating scrubs of Betadine (Purdue Products) and 70% ethanol. An incision was made along the lateral femur, and blunt dissection through the biceps femoris muscle exposed the trifurcation of the sciatic nerve into the sural, common peroneal, and tibial branches. The common peroneal and tibial nerves were isolated together, ligated tightly with 6-0 silk suture (Braintree Scientific), and transected distally, with a 1.0–1.5 mm nerve segment resected to prevent axonal regeneration. Extreme care was taken to avoid any stretch, traction, or mechanical disturbance to the adjacent sural nerve, which was left intact. The muscle was closed, and the skin incision was secured with absorbable sutures and surgical staples. Sham-operated control mice underwent identical surgical exposure of the sciatic nerve branches without ligation or transection.

### Reflexive Nociceptive Assays and c-Fos Activation

All mice undergoing behavior studies included both males and females starting at 8 weeks of age and these experiments were carried out to a maximum of 3 months. On the day of training/testing, mice were transferred to the behavior room for a 30-minute habituation.

#### Von Frey Filament Testing

Mechanical sensitivity was assessed using calibrated von Frey monofilaments (0.04 to 1.4 g; Stoelting) applied to the lateral plantar aspect of the injured (or sham) hindpaw (sural nerve dermatome)^40^ and was adapted from Murthy et al., 2018^41^. Each filament was applied 4 times perpendicularly until slight buckling occurred. The withdrawal frequency (percentage of positive paw withdrawals out of 4 stimulations) was recorded sequentially from lowest to highest force.

#### Dynamic Tactile Allodynia (Brush Test)

Dynamic brush testing was used to assay tactile sensitivity at baseline as well as dynamic allodynia during neuropathic pain^42^. Dynamic light mechanical sensitivity was measured by lightly stroking the glabrous plantar surface of the hindpaw from heel to toe with a soft paintbrush (Artlicious). Nine stimulations were delivered per session (in blocks of 3 trials separated by 5-min intervals). Responses were scored on an ordinal scale 0-3: 0=no response, 1=withdrawal, 2=guarding, 3=licking/tending. The composite brush score was expressed as the sum of all scores divided by total stimulations.

#### Mechanical Hyperalgesia (Pinprick Test)

Pinprick testing was used to assess noxious mechanical sensitivity at baseline and mechanical hyperalgesia during neuropathic pain^40^. Noxious mechanical sensitivity was probed using an insect dissecting pin (Austerlitz, Size 1) applied perpendicular to the sural territory of the plantar paw with sufficient force to indent but not penetrate the dermis. Responses were scored on an ordinal scale 0-3: 0=no response, 1=withdrawal, 2=guarding, 3=licking/tending. The composite brush score was expressed as the sum of all scores divided by total stimulations.

#### cFos Immediate-Early Gene Induction

Fourteen days post-surgery, sham and SNI mice were habituated to acrylic testing chambers for 3 consecutive days (30 min/day). On the experimental day, mice received a repetitive noxious pinprick stimulus (23-gauge BD needle) to the left hindpaw once every 60s for 30 consecutive minutes. Exactly 90 min after stimulus termination, mice were anesthetized and transcardially perfused for immunohistochemical quantification of c-Fos in the primary somatosensory cortex, anterior cingulate cortex, and DLS.

### Automated Reproducible Mechano-stimulator (ARM) Testing

To deliver standardized, operator-independent mechanical stimuli while recording high-speed behavior and fiber photometry, we deployed the Automated Reproducible Mechano-stimulator (ARM).

#### Hardware Architecture

The ARM behavior setup uses Zaber motorized linear stages for stimulus presentation^43^. Mice are situated in acrylic chambers (4.2 cm × 11.5 cm × 4.6 cm) on a specialized mesh table held down by acrylic weights. This motorized platform also includes two 500 fps cameras on each side of the chamber (0.4 MP, 522 FPS, Sony IMX287). A bottom camera allows for visualization of stimulus delivery and recordings from all 3 cameras per trial are stored.

#### Force Delivery and Closed-Loop Feedback

Stimuli were delivered using an interchangeable probe (blunt probe or pinprick) coupled to a digital load-cell force sensor (DSCUSB). The probe moved vertically through the mesh floor to depress the sural receptive field until a programmable cutoff force was reached (20g maximum threshold), maintaining the displacement until withdrawal or time elapsed. Force calibration was performed using 2g and 20g standard analytical weights.

#### Photometry Synchronization

Probe-to-paw contact was detected closed-loop via the calibrated load cell, which simultaneously triggered an Arduino Uno Rev 3 microcontroller to output a hardware TTL pulse to the Tucker-Davis Technologies (TDT) acquisition rack, synchronizing stimulus onset (t=0 s) with optical signals.

#### Testing

Mice are habituated 3x for 1 hour/day, one week before testing. The habituation program allows for the motorized stimulus to move randomly and give empty air to acclimate mice to the noise. On the day of testing, only one mouse is tested at a time and the left hindpaw is stimulated with either a blunt probe or a pinprick. Each mouse was tested for 30 trials with 1 minute rest between trials. The stimulus was delivered to the sural nerve territory of the paw.

#### High-speed Kinematic Tracking (SLEAP) and Millisecond Pain Behavior Analysis (PAWS)

To assess fine-scale kinematics high-speed lateral video recordings of mechanically evoked hindpaw withdrawal were acquired at 500 frames per second (fps) using the ARM and analyzed using Social LEAP Estimates Animal Poses (SLEAP)^44^ coupled to Pain Assessment at Withdrawal Speeds (PAWS)^45^ platform. Video recordings were synchronized to stimulus delivery, with t = 0s defined by the load-cell/Arduino contact TTL signal.

#### Spatial calibration and Coordinate Transformation

Videos were spatially calibrated in Photron FASTCAM Viewer 4 (PFV4) using the known dimensions of the ARM testing chamber (length, width, height = 11.5 cm x 4.2 cm x 4.6 cm) to determine the physical distance represented by each pixel. This calibration allowed pixel-based tracking coordinates to be converted into physical distance measurements, establishing an animal-specific scaling factor (0.16 mm/pixel) and enabled quantitative comparison of metric physical distances and velocities across experimental conditions and time points.

#### Supervised Deep-Learning Pose Estimation (SLEAP)

A deep neural network was trained in SLEAP to track hindpaw kinematics during stimulus-evoked withdrawal reflex. A diverse training set comprising roughly 100 videos (300,000 total frames) was selected from male and female mice, including SNI and sham conditions across baseline, Week 1, Week 3, and Week 12 timepoints. Roughly 800 frames from these videos (6-10 per video) were manually annotated in SLEAP by labeling three anatomical landmarks on the stimulated hindpaw: the heel, center (metatarsophalangeal joint), and toe to build the prediction model. The trained neural network performed automated inference across all experimental trial videos. Videos with incomplete predictions (< 90% of frames predicted on) and low-confidence predictions (prediction score < 0.5) were flagged as tracking occlusions and reconstructed via linear interpolation. SLEAP tracking outputs were exported as H5 files and converted to CSV format using R for compatibility with PAWS.

#### Kinematic Quantification (PAWS)

The resulting coordinate data were imported into PAWS for quantitative analysis of paw movement during mechanically evoked withdrawal responses. PAWS analysis was performed at a video frame rate of 500 fps using a fixed paw-height baseline of 1.0 and a y-threshold of 0.5 for determining activity above baseline. A p-cutoff of 0.45 was applied to tracked coordinates, with a shaking threshold of 0.35, and a paw-withdrawal threshold of 0.5. Coordinate trajectories were temporally smoothed using an average smoothing filter with a 7.0 frame window. The center of the paw (metatarsophalangeal joint) was used for the primary kinematic measurements that included: maximum paw height, maximum x-velocity, maximum y-velocity, providing an objective characterization of the behavioral response beyond the presence or absence of paw withdrawal.

### Blackbox Naturalistic Behavioral Arena

Unconstrained, multi-modal naturalistic behaviors were recorded in the Blackbox One apparatus.

#### Apparatus and Sensor Systems

The Blackbox One captured videos and Fourier Transform Infrared Spectroscopy (FTIR) data for a single mouse in a 18cm x 18cm chamber with camera imaging recording of 45-90 fps with 4MP sensor^46^. Mice are allowed to roam freely inside an opaque black chamber with an inner chamber in darkness, while recorded from below using a near-infrared (NIR) enhanced HD machine vision camera. The animal’s body pose is captured with a rapid pulse of NIR light, while the plantar contact areas and weight distribution of the paws are recorded separately with a proprietary Optitouch sensor that outputs a heatmap of the mouse’s body. These data streams are combined to generate a high-speed, high-resolution video ethogram of mice. The Blackbox utilized the Palmreader software platform that uses machine learning algorithms to automate measurement of paw pressure, body pose estimation, supervised and unsupervised behaviors.

#### Testing protocol and TTL synchronization

Mice were tested across 20-min sessions (two pre-operative baseline sessions and weekly post-operative sessions spanning 12 weeks). The Blackbox hardware continuously streamed digital TTL markers to the TDT acquisition console. The first and last TTL transitions bounded the continuous video recording, establishing a common time base (t=0 s at video frame 0) to align continuous calcium recording with computer-vision kinematic tracking.

#### Software and Feature Extraction

Specific kinematic and plantar parameters extracted from Palmreader included hindpaw luminance (measured as pixel intensity, representing the sum of all pixel intensity signals induced by a given paw in each video frame via FTIR), standing hindpaw luminance (the hindpaw luminance specifically measured during periods when the animal is classified as “standing” (i.e., neither of the animal’s front paws are contacting the floor)), toe spread (measured as the distance between the distal 1st digit and the distal 5th digit of the hindpaw), hindpaw lifted time (calculated as the total duration (in seconds) when a given hindpaw is not contacting the floor), heel distance (measured as the pixel distance between the left and right heel landmarks (specifically, ankle/heel joints)), and hindpaws distance (measured as the average pixel distance between the left and right hindpaw centroids across timepoints).

### *In Vivo* Fiber Photometry Signal Acquisition

Optical data were collected using an RZ10x real-time lock-in processor controlled via Synapse software (Tucker-Davis Technologies).

#### Excitation optics

This amplifier and accompanying Synapse software was used to control a custom-built optical benchtop through drivers (LEDD1B, Thorlabs) to modulate LED signals. This optical benchtop consisted of a self-contained system of four 30 mm cage cubes with integrated filter mounts (CM1-DCH/M, Thorlabs). A 405 nm LED (M405L4, Thorlabs) and a 470 nm LED (M470L3, Thorlabs) were mounted onto the first 30 mm cage cube. The 405 nm LED was used to extract the calcium-independent isosbestic signal, and the 470 nm LED was used to acquire calcium-dependent GCaMP signals. The 405 nm isosbestic signal was modulated at 210 Hz, and the 470 nm GCaMP signal was modulated at 330 Hz.

#### Optical train

A 425 nm dichroic longpass filter (DMLP425R, Thorlabs) in the first cage cube reflected the 405 nm excitation light and permitted the 470 nm light to pass through. As the light entered the second cage cube, both excitation lights were reflected by a 495 nm dichroic longpass filter (495DCLP, 67–079, Edmund Optics) into the third cage cube. A 460/545 nm bandpass filter (69013xv2, Chroma) reflected both excitation wavelengths down to the subject via a low autofluorescence patch cable (MAF3L1, core = 400 µm, nA = 0.50, length = 1m, Thorlabs). This cable was attached onto the implanted optical cannula by a ceramic mating sleeve (Thorlabs). Isosbestic and jGCaMP8s emissions were collected via the optic cannula and passed through the 460/545 nm bandpass filter (69013xv2, Chroma) into the fourth.

#### Demodulation

Finally, the emission fluorescence passed through the detection pathway and was focused onto the internal photodetectors of the RZ10x and was digitally demodulated in real time at a sampling frequency of F_s_ =1017.25 Hz.

### Evoked Fiber Photometry Signal Processing (ARM Assay)

Raw streams from the TDT system for GCaMP and isobestic signals were processed offline in MATLAB:

1. Initial Truncation: The initial 8,000 samples (∼7.86s) were discarded to eliminate LED warm-up artifacts and transient filter responses.
2. Polynomial Detrending: Photobleaching drift was corrected by fitting a second-order polynomial to the time series of each channel (T_s_): Ŷ_channel_(t)=at^2^+bt+c. The detrended signal was defined as: Y_detrended_(t)=Yraw(t)−Ŷ_channel_(t).
3. Robust Median/MAD Z-Scoring: Signals were standardized using the Median Absolute Deviation (MAD):

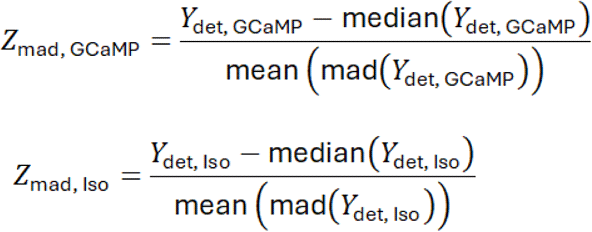
4. The motion-corrected calcium trace was obtained by subtraction: Z(t)=Z_mad, GCaMP_(t)−Z_mad_, _Iso_(t)
5. Artifact Removal: Signal excursions exceeding +4σ (high-frequency electrical spikes) or falling below −2σ (optical cable dropouts) relative to the trace standard deviation were designated as artifacts, flagged as NaNs, replaced via linear interpolation, and smoothed with a 5-point median filter.
6. Trial Segmentation: Trials were segmented from −10 s to +10 s relative to mechanical paw contact (t=0s). A non-task spontaneous control window was extracted from +10 s to +20 s post-stimulus.

#### Evoked Calcium Trace Kinetics

From the trial-averaged post-stimulus trace (t≥0 s), several parametric kinetic metrics were calculated: Peak Amplitude: maximum Z-score during post-window; Time to peak: the time required to reach peak maximum; and Decay time: the time required to reach baseline.

### Continuous Photometry Batch Preprocessing (Blackbox Naturalistic Assays)

Continuous photometry recordings during open-field Blackbox assays were processed using a batch-automated pipeline in MATLAB:

1. TTL windowing: Raw data were indexed to camera TTLs. The first rising edge was assigned as t=0 s, and all samples prior to first rising edge and after the final pulse were excised, eliminating recording initialization surges.
2. Linear Detrending and Robust Normalization: Signals were linearly detrended (detrend) to correct for slow baseline shifts. GCaMP8s and isosbestic streams were independently normalized by their median and MAD (see above). The motion-subtracted trace was computed: Z_cleaned_ = Z_GCaMP_ -Z_Iso_
3. Artifact Curation: Excursions exceeding +4σ or −2σ of the session standard deviation were marked as NaNs, linearly interpolated, and 5-point median filtered.
4. Standardized Export: Cleaned data structures containing timestamps, sampling rate (F_s_ =1017.25 Hz), normalized traces (Z_GCaMP_,Z_Iso_), and the cleaned trajectory (Z_cleaned_) were exported as .mat and .csv files for downstream behavioral alignment.

### Supervised Learning with Simple Behavioral Analysis (SimBA)

To identify classic behavioral phenotypes (grooming, rearing, locomotion), we deployed a supervised machine learning framework in the Simple Behavioral Analysis (SimBA) environment:

#### DeepLabCut Tracking

DeepLabCut (v2.2) tracked 7 anatomical landmarks from the ventral perspective: snout, neck/chest, left front paw, right front paw, left hind paw, right hind paw, and tail base exported from the Blackbox software. Frames with low confidence (p<0.1) were interpolated. Extracted coordinates were imported into the Simple Behavioral Analysis (SimBA) software framework^47^.

#### Geometric Feature Library

To overcome ventral-view occlusion challenges and accurately capture pathological pain-related guarding phenotypes, we engineered a custom 30-feature library (“Geometric Fingerprint”). Features were engineered from:

1. Inter-limb Euclidean distances: Forepaw-to-snout and hindpaw-to-snout distances.
2. Postural geometry: Snout-to-tailbase longitudinal axial length.
3. Spinal configuration: Three-point angles formed along the snout–neck–tailbase axis.
4. Animal footprint envelope: Convex hull area computed across all tracked body parts.

#### Random Forest Modeling

Classifiers were trained on 16 balanced sessions using 2,000 decision trees. Active learning (hard-negative pseudo-labeling) was implemented to differentiate pain-induced asymmetric guarding from grooming. Background frames were under-sampled (10:1 for grooming and locomotion; 20:1 for rearing) with a minimum leaf size of 5. Predictions were smoothed via Kleinberg burst detection (σ=2,γ=0.3), enforcing minimum duration thresholds: 500 ms for grooming, 1,000 ms for locomotion and rearing.

#### In Vivo Fiber Photometry Synchronization and Preprocessing

Fluorescent calcium dynamics were recorded using a Tucker-Davis Technologies (TDT) multi-channel fiber photometry system. Excitation light targeting GCaMP8s (465 nm) and an isosbestic control (405 nm) were sinusoidally modulated, focused through a patch cord, and delivered to targeted striatal subregions (D1- or A2A-expressing medium spiny neurons). Optical signals were demodulated, low-pass filtered, and detrended to correct for baseline photobleaching. Cleaned Z-scored calcium trajectories (ΔF/F normalized by baseline standard deviation) were exported alongside system timestamps (F_s_). Behavioral event timestamps (grooming, rearing, locomotion) exported from SimBA were synchronized to photometry data streams using hardware-timed Blackbox TTL triggers.

#### Behavior-Locked Calcium Dynamics (PSTH Generation)

Peri-event histograms were aligned to behavior onsets (t=0 s) across a [−2.0,+15.0]s window for bouts lasting between 1.0s and 20.0s. Local baseline

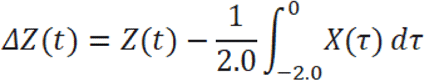

subtraction was applied using the pre-onset window:

#### State-Based auROC Analysis

Active behavioral states were defined with a +0.5s post-bout buffer ([t_start_,t_end_+0.5 s]) to accommodate GCaMP8s dissociation kinetics. Continuous discriminability was computed via Mann-Whitney U test, and statistical significance was evaluated with 10,000 circular permutations.

### Unsupervised Learning with Variational Animal Motion Embedding VAME

To quantify unbiased, naturalistic motion repertoires without predefined human classifications, continuous tracking data were processed within the Variational Animal Motion Embedding (VAME) framework^48^:

#### Egocentric Alignment and Feature Cropping

From the full 57-keypoint Blackbox DeepLabCut model, tracking coordinates were egocentrically centered on the animal’s centroid and aligned along the anteroposterior vector connecting the tail base to the snout. The coordinates were then down-selected to a 6-keypoint core kinematic set (12 coordinates: snout, left front paw, right front paw, left hind paw, right hind paw, and tail base). Time series were formatted to 12×frames and Z-scored within each session.

#### RNN-VAE Architecture and Syllable Segmentation

Kinematic patterns were embedded using a recurrent neural network (RNN) variational autoencoder (VAE) consisting of bidirectional Gated Recurrent Units (GRU) compressing temporal sequences into a continuous latent space of z=30 dimensions. The model was trained on a balanced cohort of 38 representative sessions spanning all conditions (Sham vs. SNI) and longitudinal timepoints. The trained network projected all 218 experimental sessions into the common latent manifold. An autoregressive Hidden Markov Model (HMM) segmented continuous trajectories into 15 discrete behavioral syllables (K=15).

#### Macro-Scale Behavioral Communities and Grammar

To capture ethologically relevant behavioral macro-states that align with physiological calcium signaling kinetics, the 15 motifs were organized into higher-order “Communities” through hierarchical agglomerative clustering:

1. Global Transition Probability Matrix: For each session, a first-order Markovian transition probability matrix *P* was calculated where element *P_ij_* represents the conditional probability of transitioning from motif *i* to motif *j*

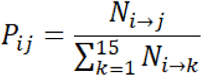

The mean transition matrix was computed across all 218 sessions.

2. Community Partitioning: A pairwise distance matrix was defined based on bidirectional transition affinities:

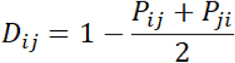

with diagonal elements set to zero. Agglomerative clustering was executed using Ward’s minimum variance method. Cutting the resulting dendrogram at an empirical distance threshold of 1.0 partitioned the 15 motifs into five distinct behavioral communities.

3. Biological Validation: Community identities were visually ground-truthed by generating video montages of representative bouts across 14 benchmark sessions. To correct for the temporal convolution and sliding window latency inherent to the VAME RNN architecture, video extraction incorporated a +22 frame offset (488.9 ms) between VAME temporal indices and source video frames. Montages defined the communities as:

- Community 1: Low-speed Locomotion (Motifs 0, 9)
- Community 2: Quiescence / Resting (Motifs 5, 10, 12)
- Community 3: Micromovements / Fine-Motor Investigation (Motifs 6, 8, 14)
- Community 4: Exploratory Movement (Motifs 1, 2, 4)
- Community 5: High-speed Locomotion (Motifs 3, 7, 11, 13)

Macro-Grammar Dynamics and Transition Statistics: For each session, macro-state dynamics were quantified across several metrics:

1. Community Occupancy: The proportion of total session duration spent within a given community:

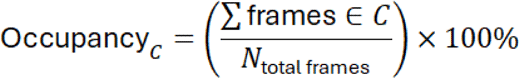
2. Mean Bout Duration: Continuous bouts of a community were identified by changes in state labels (Δlabel ≠ 0). The mean duration (in seconds) of contiguous frames spent in state *C* before transitioning was calculated as:

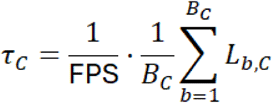

where *L_b,c_* is the frame length of bout *b*, and *B_c_* is the total number of bouts of community *C*.

3. Behavioral Transition Entropy: To quantify the stereotypic predictability versus stochastic variability of behavioral sequencing, Shannon entropy (*H*) was calculated across the rows of the 5×5 community transition probability matrix (*P_comm_*):

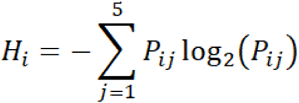

Global behavioral entropy was defined as the mean entropy across all active states 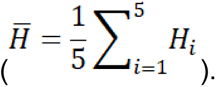. Lower entropy denotes restricted, stereotyped transitions, whereas higher entropy indicates flexible, exploratory behavioral syntax.

4. Transition Pair Significance (Chi-Square Analysis): Direct comparisons of individual transition probabilities between Sham and SNI at specific timepoints (e.g., Week 1, Week 3, Week 12) were computed from contingency tables of observed transition counts (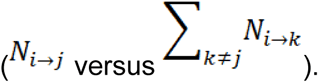). Laplace smoothing (+1 pseudocount) was applied to avoid division-by-zero artifacts for low-probability transitions. Statistical divergence was evaluated using two-tailed Pearson’s Chi-square contingency tests (*χ*^2^).

#### Longitudinal Behavioral Grammar Trajectory and Abnormality Index

To map the longitudinal adaptation of behavioral structure following nerve injury:

1. UMAP Embedding: The 5×5 transition probability matrix of each session was flattened into a 25-dimensional feature vector *T*^→^ = [*T*_1→1_, *T*_1→2_,…, *T*_5→5_]. Feature vectors across all 218 sessions were standardized (Z-score normalization) and projected into a low-dimensional manifold using Uniform Manifold Approximation and Projection (UMAP; parameters: n_neighbors=15, min_dist=0.1, Euclidean metric, random_state=42).
2. Centroid Permutation Testing: At each longitudinal timepoint, the observed behavioral separation between Sham and SNI cohorts in UMAP space was quantified by the Euclidean distance between group centroids:

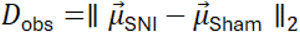

To assess significance, condition labels were randomly permuted 1,000 times, recalculating centroid separation to generate a null distribution (*D_nul_*_l_). The empirical *p*-value was defined as:

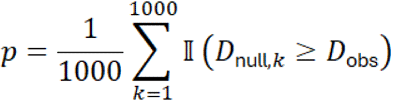

3. Mahalanobis Pain/Abnormality Index: To establish a continuous subject-level readout of behavioral divergence, we calculated the Mahalanobis distance (D_M_) of each individual mouse’s 2D UMAP coordinate relative to the multivariate normal distribution of the healthy control (Sham) cohort at that same timepoint:

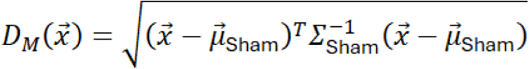

where, *x*^→^ [*U*, *MAP*_1_, *UMAP*_2_], *μ*^→^_sham_ is the mean vector of the Sham group, and *Σ*_sham_ is the covariance matrix of the Sham group. This distance quantifies behavioral pathoneurotic drift in units of standard deviations from healthy baseline variability.

### Community-Locked Photometry and auROC Discriminability

Cleaned continuous photometry streams (*F_s_* _=_ 1017.25 Hz) were aligned to VAME community time series (45fps):

1. Temporal Synchronization: The alignment ratio was defined as

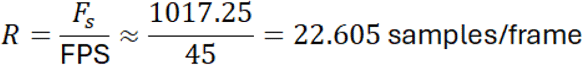

Community onset timestamps incorporated the +22-frame model latency correction:

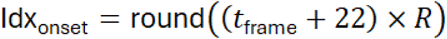
2. Bout Duration Filter: Only community bouts persisting for ≥1.0 s (≥45 consecutive video frames) were included.
3. PSTH Snippet Extraction: Calcium snippets were extracted from −2.0s to +8.0s relative to community onset and baseline-subtracted using the pre-onset window (−2.0s to 0s).
4. Non-Parametric Area Under the ROC Curve (auROC) State Discriminability: A continuous binary mask *M(t)* was generated at 1017.25 Hz across the entire session (M=1 during execution of the target community; M=0 during all other behaviors). Calcium distributions during the behavior (*Z*_in_=*Z*[*M*=1]) and outside the behavior (*Z*_out_=*Z*[*M*=0]) were compared via the Wilcoxon rank-sum test. The Mann-Whitney U statistics were calculated as:

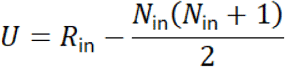

where R_in_ is the rank sum of the in-state samples. Empirical discriminability was computed as:

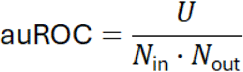
5. Circular Permutation Shuffling: To calculate significance while preserving signal autocorrelation, the state mask M(t) was circularly shifted 10,000 times by a random offset Δ*t_s_* ∊ [*τ*_min_, T - *τ*_min_] where the minimum shift buffer was set to *τ*_min_= 20 s(20,345 samples), and *T* was total session length. A two-tailed empirical p-value was derived:

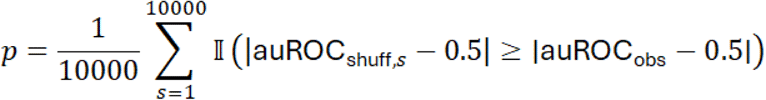
6. Population Statistics: Group distribution shifts between Baseline 2 and Week 3 were tested using the two-sample Kolmogorov-Smirnov test, two-sample F-tests for equal variances, and bias-corrected Hedges’ g:

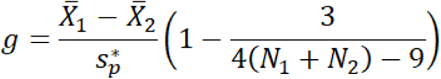

### Acute Brain Slice Preparation and Whole-Cell Patch-Clamp Electrophysiology

Mice were briefly induced (via 3% isoflurane), deeply anesthetized with an intraperitoneal injection of ketamine-xylazine (300/30 mg/kg, respectively), and transcardially perfused with recovery artificial cerebrospinal fluid (ACSF), which contains the following (in mM): 103 NMDG, 2.5 KCl, 1.2 NaH_2_PO_4_, 30 NaHCO_3_, 20 HEPES, 25 Glucose, 101 (1 N) HCl, 10 MgSO_4_, 2 Thiourea, 3 Sodium Pyruvate, 12 N-Acetyl-L-Cysteine, and 0.5 CaCl_2_ (saturated with 95% O2 and 5% CO2^49^. Following decapitation, the brain was rapidly extracted and submerged in recovery ACSF before being mounted onto a VT-1200S vibratome (Leica). The vibratome chamber was filled with oxygenated recovery ACSF, and 300 µm slices were cut. Once the hippocampus began to appear, vibratome sectioning was terminated, and the posterior tissue block containing the injection site was transferred into 10% neutral-buffered formalin for post-hoc confirmation. Slices were immediately transferred to recovery ACSF that was heated to 35 °C for ∼5 min. After, slices were transferred to room temperature, external ACSF which contained (in mM): 124 NaCl, 2.5 KCl, 26 NaHCO_3_, 1.2 NaH_2_PO_4_, 10 Glucose, 3 Sodium Pyruvate, 1 MgCl_2_, and 2 CaCl_2_ (saturated with 95% O2 and 5% CO2). They were allowed to recover for at least ∼45 min before recording. Whole-cell patch clamp recordings were acquired from slices that were constantly superfused (2-4 mL/min) with oxygenated external ACSF at ∼34 °C. Slices and cells were visualized by infrared differential interference contrast (IR-DIC) microscopy using a CCD camera (Hamamatsu) mounted onto a BX-51WI upright microscope (Olympus) fitted with a swinging objective holder containing two switchable lenses: a 4x air lens and a 40x water-immersion lens. Patch pipettes (3-5 MΩ) were pulled from borosilicate glass micropipettes (2 mm O.D., Warner Instruments) using a P-1000 horizontal puller (Sutter Instruments).

#### Electrophysiology Feature Extraction

Current-clamp recordings were obtained from unidentified and identified striatal neurons in mice expressing tdTomato in either D1-SPNs or D2-SPNs within anterior DLS (AP from bregma: 0.5 to -0.7mm) which is known to receive S1 inputs^6^. The internal pipette solution for current-clamp experiments contained (in mM): 130 K Methanesulfonate, 10 KCl, 10 HEPES, 2 MgCl_2_, 4 Na_2_ATP, 0.4 Na_2_GTP, at pH 7.25 and 285–290 mOsm/L. ChR2 in the S1HL axon terminals was stimulated via illumination with a 2.5ms, 470 nm LED light pulse (∼0.6 mW measured after the objective; Thorlabs) delivered through the 40x objective lens. The illumination spot size had a diameter of ∼550 µm. Once a stable gigaseal was formed between the recording pipette and cell membrane, negative pressure was applied to rupture cell membrane to gain access to the cell interior. The internal solution was permitted to dialyze for ∼5 min. At this point, access resistance was monitored and the resting membrane potential was recorded. All cells were held at –80±2 mV to ensure equal driving forces when studying synaptic strength and short-term synaptic plasticity. Baseline voltages that drifted outside of this range were excluded from analysis. Patched cells were held for ∼25-35 min while being run through a standardized set of protocols: 1) hyperpolarizing/depolarizing current steps (–300 pA to +400 pA, 50 pA steps, 500ms, 15 sweeps) to evaluate cell health and intrinsic parameters, 2) single pulse (single 2.5ms blue light pulse was presented once every 15 s for 20 sweeps), 3) paired pulse ratio (five 2.5ms pulses, separated by 50ms inter-pulse intervals (IPI), were presented once every 15 s for 20 sweeps), and 4) train stimulation (thirty 2.5ms pulses, separated by 64.2ms IPI (15 Hz), were presented once every 30s for five sweeps). While these protocols were being run, biocytin within the internal solution diffused into the cell for subsequent 3D morphological reconstructions. Occasionally, unidentified cells within the same field of view were sequentially patched following the initial identified cell patch to control for injection site volume and location. Data was acquired via a EPC10USB amplifier and digitized at 20 kHz in Patchmaster Next (HEKA). Liquid junction potential was not corrected in these traces.

#### Analysis of patch clamp recordings

All data were analyzed offline using custom-written MATLAB and Python scripts. DAT files from Patchmaster Next (HEKA) were imported, organized, and saved as a .mat file. The .mat file data was imported into Python for post-processing using the electrophysiology feature extraction librarIFEL created by the Blue Brain Project^38, 50^. Intrinsic parameters were calculated at the +350 pA current step. Key values pertaining to every action potential in a sweep were calculated including the action potential threshold value and index, the peak value and index, and the corresponding minimum afterhyperpolarization (AHP) value and index. Each action potential threshold value and index were captured using a derivative threshold method (dV/dt ≥10 mV/s). Action potential properties were assessed for all spikes in a sweep and averaged together. The action potential peak was defined as the difference between the action potential threshold and its maximum positive peak. Half-height width (HHW) and rise time were both calculated via interpolation. HHW was measured at 50% of the action potential peak, while rise time was measured from 10% to 90% of the action potential peak. Rheobase was calculated as the first current step which elicited action potentials. AHP amplitude was calculated as the difference between the action potential threshold value and the minimum AHP value. For all optogenetically evoked PSPs, a baseline measure was calculated by averaging the first 10,000 sampling points for each individual sweep. This measured value was subtracted from every value in each individual sweep, resetting the baseline equal to zero. The maximum PSP amplitude, relative to the zero-baseline value, was calculated from an average of 20 sweeps during single pulse and paired-pulse protocols, and 5 sweeps during train protocols. The time to maximum PSP amplitude was measured as the difference between photostimulation onset and the maximum PSP index.

### Histology, Immunohistochemistry, and Confocal Microscopy

#### Histology

Mice were anesthetized with an intraperitoneal injection of Ketamine-Xylazine (120 mg/kg Ketamine, 24 mg/kg Xylazine) and transcardially perfused with PBS followed by 10% neutral-buffered formalin. The brain was extracted and stored in 10% neutral-buffered formalin overnight at 4°C. The following day, they were transferred into 0.2% PBS Azide at 4°C. Tissue was mounted onto a stage and sectioned at 100 µm using a VT-1000 vibratome (Leica). Slices were mounted onto microscope slides using DAPI Fluoromount-G (Southern Biotech #0100–20) and coverslipped before confocal imaging. For all behavioral experiments, injection site location verification, cannula placement, and viral expression in the DLS were confirmed. For electrophysiology experiments, slices from recordings containing the injection site were processed.

#### cFos Immunostaining

For cfos experiments, mice were anesthetized (5% induction, 1% maintenance) and transcardially perfused with PBS followed by 10% neutral-buffered formalin. Brains were stored in formalin for 3 hours and washed 3x with PBS Azide. Coronal sections of the dorsal striatum, anterior cingulate cortex, and S1HL (∼50um) were taken and stained with primary antibody, 1:1000 rabbit anti-cFos (Synaptic systems, 226008), and then with secondary antibody, 1:1000 Goat anti-Rabbit IgG (H+L) Alexa Fluor 594 secondary antibody (Life Technologies, A11037), and with 1:500 DAPI.

### Quantification and General Statistical Analyses

Data were compiled and analyzed using GraphPad Prism (version 9.0), MATLAB (R2022b), and Python (3.9+). Graph Pad (version 9.0). Normality was assessed using the Shapiro-Wilk test. Continuous parametric datasets with repeated measurements across timepoints or stimuli were evaluated via unpaired Student’s t-tests, two-way repeated-measures ANOVA, or mixed-effects models (restricted maximum likelihood, REML) when values were missing. Significant main effects or interactions (F-test p<0.05) were evaluated with Sidak’s or Tukey’s post-hoc tests with adjustment for multiple comparisons. Ordinal data, non-normally distributed parameters, and categorical behavioral scores were evaluated using the Mann-Whitney U test. Transition probabilities and contingency tables were tested via two-tailed Pearson’s *X*^2^ tests with Laplace continuity smoothing. Centroid UMAP separation and state-based auROC discriminability metrics were tested via non-parametric permutation tests (N=1,000 and N=10,000 iterations, respectively). Statistical significance was defined at α=0.05, denoted as:∗p≤0.05,∗∗p≤0.01,∗∗∗p≤0.001,∗∗∗∗p≤0.0001. All summary data are presented as mean ± standard error of the mean (s.e.m.) or median with interquartile range (IQR), as indicated in the figure legends. Exact statistical tests, degrees of freedom, sample sizes (n representing either mice, sessions, or cells), and effect sizes (e.g., Hedges’ g) are reported in each figure legend.

## Results

### Evoked stimulus response differentially engages striatal SPN populations during neuropathic pain

Nociceptive information is highly organized within the spinal cord with the dorsal horn acting as the primary central relay and modulation site originating from the periphery. Pain information travels through ascending projections via the spinothalamic tract and terminates in the primary somatosensory cortex (S1)^7^. S1 and many other brain regions related to pain function send projections to the striatum^6^ and notably, neuropathic pain induces hyperactivity in S1^8–10, 51^, making this corticostriatal pathway a critical area to study underlying circuit changes during pain-related behavior. Therefore, our focus centered on downstream projections from S1, namely the dorsal striatum (DStr) that could potentially have a role in coordination of sensory input and motor outcomes. Accordingly, we injected an anterograde tracer, pAAV-CAG-tdTomato, into the hindlimb region of S1 (S1HL) which revealed dense projections to the dorsolateral striatum (DLS) (**Supplementary Fig. 1D**). Considering the DStr is a sensorimotor hub and there is some evidence suggesting the role of the DStr in the pain response, we further explored if the DStr is active during pain-related behaviors. We used the spared nerve injury (SNI) as our pain model given that it allows for consistent pain behavioral testing with high reproducibility to the non-injured, intact sural nerve adjacent to the transected tibial and peroneal nerves, without affecting daily life activities^52^. To measure neural activation after evoked stimulation, we quantified cFos+ cells after after 30 minutes of evoked pinprick stimulation in DLS and DLS-projecting S1HL neurons (using a retrograde tracer in the DLS, pAAVrg-CAG-GFP). We found that in the neuropathic pain condition, both the DLS and S1HL had higher cFos+ expression compared to sham controls (**Fig.1A**). We observed activation of ACC in both conditions, which may reflect an acute response to the pinprick test stimulus. Thus, the S1HL corticostriatal pathway is activated during chronic pain conditions.

**Figure 1:**
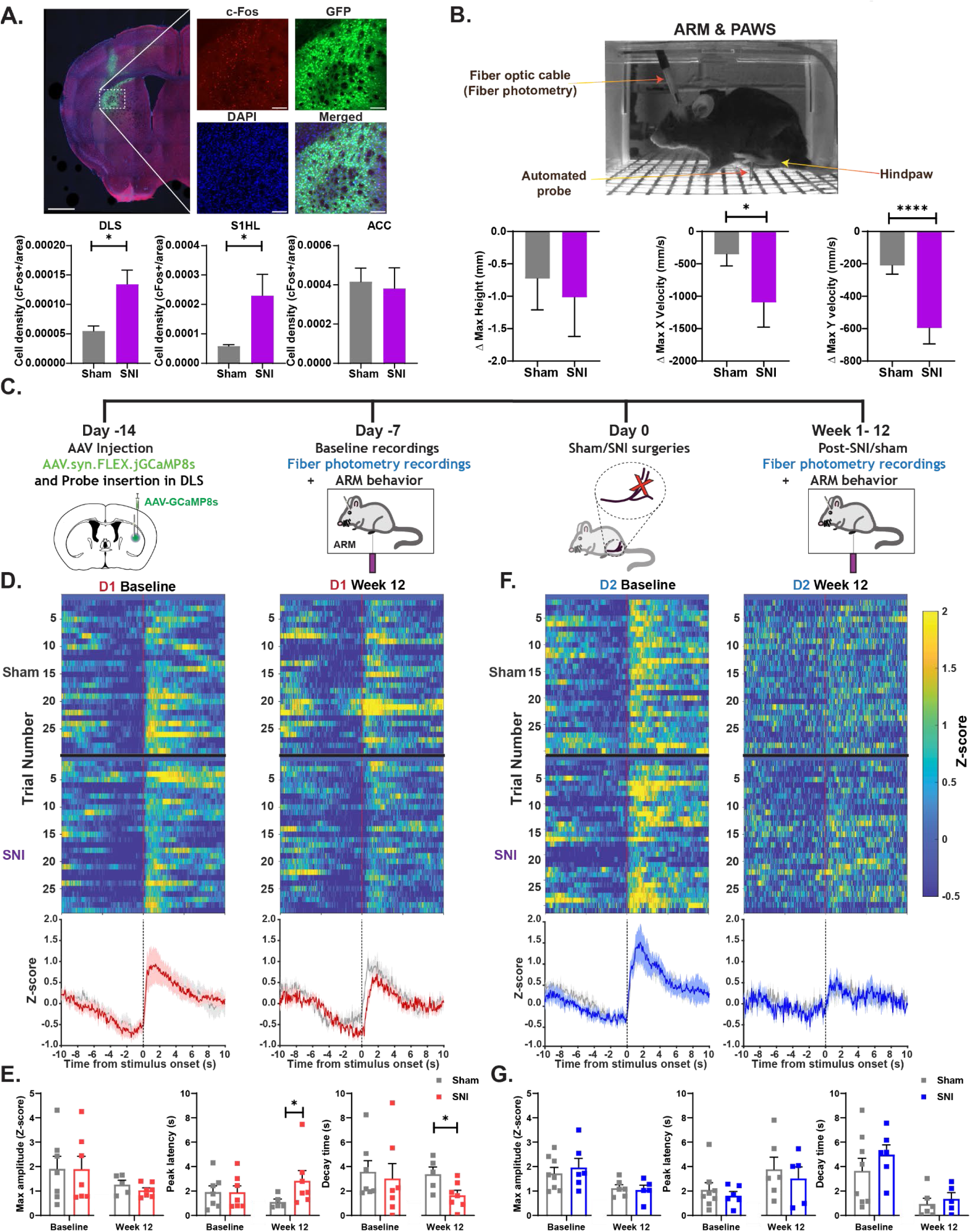
Neuropathic pain induces behavioral changes corresponding to SPN population changes. **(A)** (*Top*) Image captured from ARM video illustrating mouse in ARM chamber with fiber optic attachment and automated probe during evoked blunt stimulation. (*Bottom*) Maximum height, maximum x velocity, and maximum y-velocity evaluated by baseline subtraction (Δ = 12 Weeks - Baseline). Significant differences were observed for maximum x velocity (*p = 0.0454) and maximum y velocity (****p<0.0001), but not maximum height (p = 0.3573). Mann-Whitney U. N = 11 for sham and 12 for SNI. Error bars represent ± SEM. **(B)** Immunohistological staining of cFos (red), retro-GFP (green), and DAPI (blue) after mice were poked with a pinprick stimulus on the ipsilateral (”injured”) paw 14-days post-SNI. Image of dorsolateral striatum (DLS) is taken at 10x, inset images taken at 20x. Scale bar is 1000μm, inset 100μm. (Bottom) Quantification of cFos+ cells within GFP expression in the DLS, primary somatosensory cortex, hindlimb (S1HL), and anterior cingulate cortex (ACC). Significant differences occurred in the DLS (p = 0.0433) and S1HL (p = 0.0325), but not ACC (p = 0.1251). Unpaired t-test. N = 3 per condition. Error bars represent ± SEM. **(C)** Timeline depicting AAV-GCaMP viral injection, pre-injury fiber photometry recording, and post-injury photometry recording during evoked stimulus ARM experiments. **(D)** (*Top*) Heatmaps averaging all D1-Cre animals for sham and SNI across 30 trials at Week 12. (*Bottom*) Mean traces depicting D1 activity during evoked stimulation in the ARM apparatus aligned to probe contact with the paw. Shaded region represents SEM. **(E)** Quantification of z-scored GCaMP8s in D1-Cre mice showing significant differences in time to peak (*p = 0.0278) and decay time (*p = 0.0240), but not maximum amplitude (p = 0.2404) at Week 12. There were no differences during baseline. Mann-Whitney U. N = 5 for sham and N = 7 for SNI. **(F)** (*Top*) Heatmaps averaging all A2a-Cre animals for sham and SNI across 30 trials at Week 12. (*Bottom*) Mean traces depicting D2 activity during evoked stimulation in the ARM apparatus aligned to probe contact with the paw. Shaded region represents SEM. **(G)** Quantification of z-scored GCaMP8 in A2a-Cre mice showing no differences in D2 activity between sham and SNI mice at Week 12 or baseline. N = 6 for sham and N = 5 for SNI.

SNI has a robust phenotype that manifests into rapid (<24hr) and long-lasting (>6 months) mechanical allodynia^39^, representing a model that can be used to study chronic pain timescales relevant to clinical outcomes. To establish how SNI confers changes in motor output on a longer clinically relevant time course, we first validated hypersensitivity to mechanical stimuli in the SNI model. We did pre- and post-assessments of the ipsilateral, affected hindpaw via manual stimulation with Von Frey (VF), brush, and pinprick stimuli to evaluate pain sensitivity. We found that at the 0.04g VF probe, SNI showed increased withdrawal responses consistently at 7 days, 25 days, and 12 weeks (****p<0.0001) post-injury compared to sham controls (**Supplementary Fig. 1A)**. Some withdrawal response differences disappeared as the VF gradient increased at 1.4g, but still apparent at 12 weeks. The same animals were tested with brush and pinprick stimulation with scores of 0 = no response, 1 = withdrawal, 2 = guarding, 3 = licking/tending^53^. For both brush and pinprick stimulation, SNI animals demonstrated higher brush and pinprick scores reflecting not only increased withdrawal responses, but also higher frequency of guarding and licking suggestive of changes in affective pain features.

While our manual scoring revealed changes in affective pain features, we sought to further explore sensory-reflexive features of pain by utilizing the automated reproducible mechanostimulator (ARM)^43^, which consistently delivers an automated stimulus with the same force during each trial on a motorized platform and reduces experimenter bias seen with manual scoring. We used a blunt probe to target the ipsilateral, affected paw for 30 trials each session, over a period of 12 weeks. Applying Social Estimates Animal Poses (SLEAP)^44^ to high-speed ARM videos, we tracked the trajectory of the hindlimb and quantified sensory reflexive behavioral features using Pain Assessment at Withdrawal Speeds (PAWS)^45^. At week 1, when normalized to baseline (Δ = post-injury - baseline), we found that SNI exhibited significantly greater reductions (larger negative shifts) in maximum height (*p = 0.0459), maximum x velocity (*p = 0.0360) and maximum y velocity (****p<0.0001) compared to sham controls (**Supplementary Fig. 1B**), indicating acute pain-induced suppression of motor vigor. At week 3, group differences persisted in maximum y velocity (**p = 0.0031) but were no longer significant for maximum height or x velocity (**Supplementary Fig. 1C**). By 12 weeks post-injury, SNI animals continued to display pronounced kinematic alterations in the injured hindlimb, characterized by significantly greater reductions in maximum x (*p = 0.0454) and maximum y (****p<0.0001) velocities relative to shams (**Fig.1A**), though maximum height values converged back toward baseline values. Although an exaggerated withdrawal speed is often expected, the long-term progression of this neuropathic model is characterized by mechanical constraints suggesting that the mechanical hypersensitivity becomes uncoupled from active withdrawal vigor.

To understand how this uncoupling of mechanical hypersensitivity and withdrawal vigor is encoded in the striatum, we measured D1 and D2 SPN activity during this automated evoked behavior response. Both preclinical and clinical evidence point to specific striatal cell types that mediate pain modulation and analgesia, notably dopamine receptor 2 (D2) cells. Hence, we hypothesized that neuropathic pain modulates spiny projection neuron (SPN) population activity within the DLS that changes over time in response to evoked stimuli. For this reason, we measured bulk calcium activity via fiber photometry in both D1- and D2-SPNs by injecting both the cre-dependent genetically encoded calcium indicator, pGP-AAV-syn-FLEX-jGCaMP8s, and a cannula into the DLS of D1-Cre and A2a-Cre (D2) mice (**Fig. 1C**). There were no differences between sham and SNI for either D1 or D2 activity at baseline (**Fig. 1E,G**). However, at week 1 post-SNI, both D1- and D2-SPNs begin to show changes in temporal dynamics with both having a longer time to peak in the pain condition compared to sham controls (**Supplementary Fig. 1E**). At a typical chronic pain timepoint for preclinical studies at Week 3 post-SNI, D1-SPNs still persistently show a longer time to peak while D2-SPNs no longer show any differences in amplitude or timing dynamics (**Supplementary Fig. 1F**). Extending the study at 12 weeks post-SNI, D1-SPNs still have a longer time to peak and slower decay times compared to sham controls, without any differences in maximum amplitude (**Fig. 1E**). There were still no differences within D2-SPNs in the pain condition (**Fig. 1G**). This data highlights that although the pain condition does not change the magnitude of SPN activity, the timing of D1- and D2-SPN activity are particularly important during the acute stage of pain. Only D1-SPNs show consistent timing changes well into the late chronic phases of pain, suggesting that persistent alterations in D1-SPN timing underlie the mechanical constraints observed in late chronic time points.

### Neuropathic pain promotes microscale and complex behavioral alterations

We established neuropathic pain promotes behavioral and neural changes in the DLS during late chronic timepoints in response to an evoked stimulus. The DStr is vital for selection, initiation, and refinement of motor actions, and therefore, a critical candidate for mediating pain-induced behavioral adaptations. To understand the full repertoire of movement changes during neuropathic pain beyond evoked stimulus pain response, we examined naturalistic behaviors using the same animals from our ARM studies. These animals participated in an open field paradigm using the Blackbox One machine learning apparatus^46^, which automates measurement of paw luminance and employs its own pose estimation model using DeepLabCut to capture each animal’s movement (**Fig. 2A**). Overall, there were no differences in distance traveled, overall luminance, or pressure index (**Supplementary Fig. 2A**). Nevertheless, SNI animals demonstrated severe changes in the injured limb over the course of 12 weeks including hindpaw pressure, standing, toe spread, and lifting the paw (**Fig. 2B-E**). In fact, SNI animals also showed abnormalities in kinematic features of heel distance (main effect of condition, ***p = 0.0008) and hind paw distance (main effect of condition, ****p<0.0001). These specific shifts in gait and posture dynamically varied across acute (week 1) and late chronic (Week 12) stages of pain (**Fig. 2F**).

**Figure 2:**
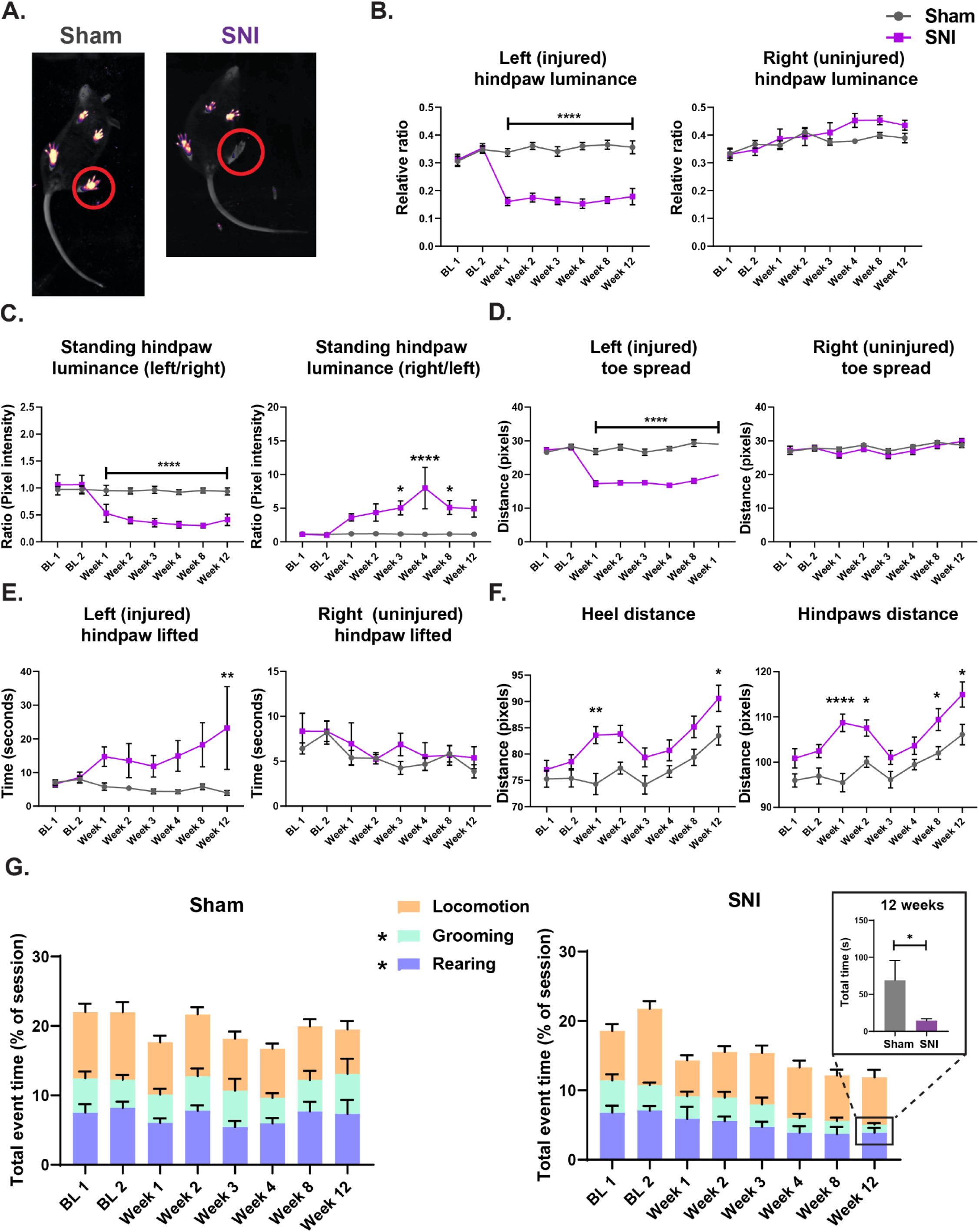
Neuropathic model demonstrates changes in paw dynamics and naturalistic behaviors. **(A)** Images captured from recordings of the Blackbox One. Red circle depicts the uninjured and injured left paw showing differences in pixel illumination. **(B)** (*Left*) Quantification of left injured hindpaw luminance over the course of baseline (BL) and weeks 1-12. There was an interaction effect (****p <0.0001), main effect of condition (****p<0.0001) and time (****p<0.0001) with significant differences at weeks 1-12 (****p<0.0001). (*Right*) Quantification of right uninjured hindpaw luminance during the same time course. There were no significant differences. Mixed-effects model analysis with Sidak’s multiple comparisons. N = 15 for sham and N = 13 for SNI. Error bars represent +/- SEM. **(C)** (*Left*) Quantification of standing hindpaw luminance (ratio of left paw/right paw) over the course of baseline (BL) and weeks 1-12. There was an interaction effect (****p <0.0001), main effect of condition (****p<0.0001) and time (****p<0.0001) with significant differences at week 1(*p = 0.0108), week 2 (***p =0.0003), week 3, 4, 8 (****p<0.0001), and week 12 (**p = 0.0041). (*Right*) Quantification of standing hindpaw luminance (ratio of right paw/left paw) during the same time course. There was an interaction effect (***p = 0.0003), main effect of condition (***p = 0.0004) and time (p = 0.0004) with significant differences at week 3 (*p = 0.0226), week 4 (****p<0.0001), week 8 (*p = 0.0232). Mixed-effects model analysis with Sidak’s multiple comparisons. N = 15 for sham and N = 13 for SNI. Error bars represent +/- SEM. **(D)** (*Left*) Quantification of left injured hindpaw toe spread over the course of baseline (BL) and weeks 1-12. There was an interaction effect (****p <0.0001), main effect of condition (****p<0.0001) and time (****p<0.0001) with significant differences at weeks 1-12 (****p<0.0001). (*Right*) Quantification of right uninjured hindpaw toe spread during the same time course. There were no significant differences. Mixed-effects model analysis with Sidak’s multiple comparisons. N = 15 for sham and N = 13 for SNI. Error bars represent +/- SEM. **(E)** (*Left*) Quantification of time spent lifting left injured hindpaw over the course of baseline (BL) and weeks 1-12. There was a main effect of condition (*p = 0.0119) with significant differences at week 12 (**p = 0.0061). (*Right*) Quantification of time spent lifting the right uninjured hindpaw during the same time course. There were no significant differences. Mixed-effects model analysis with Sidak’s multiple comparisons. N = 15 for sham and N = 13 for SNI. Error bars represent +/- SEM. **(F)** (*Left*) Quantification of distance between hindpaw heels over the course of baseline (BL) and weeks 1-12. There were main effects of condition (***p = 0.0008) and time (****p<0.0001) with significant differences at week 1 (**p = 0.0011) and week 12 (*p = 0.0430). (*Right*) Quantification of distance between the hindpaws during the same time course. There were main effects of condition (***p = 0.0008) and time (****p<0.0001) with significant differences at week 1 (**p = 0.0011) and week 12 (*p = 0.0430). Mixed-effects model analysis with Sidak’s multiple comparisons. N = 15 for sham and N = 13 for SNI. Error bars represent +/- SEM **(G)** Quantification percentage of time spent grooming, rearing, and locomotion over the course of baseline to week 12 post-injury. For grooming, there was a main effect of condition (*p = 0.0339) with significant differences specifically at week 12 (*p = 0.0281). For rearing, there was a main effect of condition (*p = 0.0295) without post-hoc differences. There were no significant differences in time spent locomotion. Mixed-effects model analysis with Sidak’s multiple comparisons. Inset, SNI spent significantly less time grooming compared to sham (Unpaired t-test, *p = 0.0439). N = 15 for sham and N = 13 for SNI. Error bars represent +/- SEM.

Building on these persistent, microscale shifts in posture and gait, alongside the delayed temporal dynamics observed in D1-SPNs during evoked responses, we reasoned that this combined mechanical and neural disruption would impair the execution of continuous motor sequences, manifesting in broader, more complex behavioral adaptations over time. We implemented supervised learning using Simple Behavioral Analysis (SimBA)^47^ and analyzed three types of behaviors, leveraging the DeepLabCut model provided from our Blackbox studies: grooming, rearing, and locomotion. Grooming is a stereotyped, self-soothing, maintenance behavior that was classified as face washing and body cleaning. Rearing was classified as mice stood on both hindpaws including up against the wall of the apparatus. Locomotion was classified as standard walking in which the mouse had to take at least 2 steps. Within all three behaviors, SNI mice spent less percentage of time grooming and rearing without any differences in locomotion (**Fig. 2G; Supplementary Fig.2B**). Specifically, SNI animals spend less time grooming at 12 weeks post-injury (**Fig. 2G, insert**). This data indicates that the pain condition produces both persistent changes in paw dynamics evident at much later chronic stages and leads to maladaptive changes in overall, complex behaviors.

### Altered naturalistic behavior is correlated with D1 and D2 SPN dynamic changes

Having observed that grooming and rearing behaviors selectively diminish during late chronic pain (Week 12) and given that the DLS is a well-established mediator of grooming^54–56^, we hypothesized that neuropathic pain alters how DLS SPNs encode these complex behaviors over time. Using the same animals from our ARM fiber photometry studies, we measured calcium activity of D1- and D2-SPNs during naturalistic behaviors in the Blackbox apparatus, up to 12 weeks (**Fig. 3A**) and correlated D1- and D2-SPN activity with grooming (**Fig. 3B**) and rearing behavior (**Fig. 3E**). We calculated the Area Under the Receiving Operating Characteristic (auROC) curve for each animal and each session and compared the distribution of Z-scored calcium signals during grooming behavior against the distribution of signals during all other times (non-behavior). An auROC is a metric of discriminability in which 0.5 reflects the calcium signal is indistinguishable from the signal from chance. An auROC of 1.0 represents perfect positive modulation, where the calcium fluorescence is consistently higher during every moment of behavior than during the background and an auROC of 0 represents a perfect negative modulation or suppression, where fluorescence is consistently lower during the behavior than during background periods. In the pain condition, D1-SPNs transitioned from a neutral state at baseline to a state of significant positive modulation during the remaining grooming bouts at week 12 (**p = 0.0087), demonstrating an increase in their association with grooming behavior (**Fig. 3C**) without any changes in sham controls. Conversely, while D2-SPNs are normally suppressed during baseline grooming, a Week 12 trend toward reduced suppression in SNI animals (*g* = -0.84) (**Fig. 3D**) was mirrored in sham controls suggesting a time-dependent effect. During rearing behavior, we see a similar effect in the D1-SPN population. In SNI animals, D1-SPNs not only shifted to a significant positive modulation at week 12 (KS, *p = 0.027), they are also hyper-associated with rearing (*g* = -1.88) (**Fig. 3F**), twice as strong as the effect with grooming. Although in the sham controls there are no significant differences in distribution between baseline and week 12, there is a large effect size (*g* = -0.7977) suggesting there is a natural drift in the circuit over 12 weeks. The D2-SPN population remains unchanged (**Fig. 3G**). We also correlated D1- and D2-SPN activity with locomotion (**Supplementary Fig. 3B**) and found that D1-SPNs of both sham and SNI conditions shift towards increased association with locomotion as time progresses (**Supplementary Fig. 3C**), indicating that the shift in locomotion is not uniquely caused by pain. However, the variance was significant in the pain condition (*p = 0.0197) revealing that while the sham group drifts upward together in a tight cluster, the SNI group is more scattered, implying that chronic pain introduces heterogeneity in the D1 population. D2-SPNs in the pain condition also show a non-significant shift towards a decreased association with locomotion (*g* = 0.8605) while the sham condition remains stable (**Supplementary Fig. 3D**). As a whole, rather than promoting an overall shift in how SPNs modulate general locomotion, persistent pain drives a profound functional reorganization within the D1 population around complex behaviors like grooming and rearing during the late chronic state.

**Figure 3:**
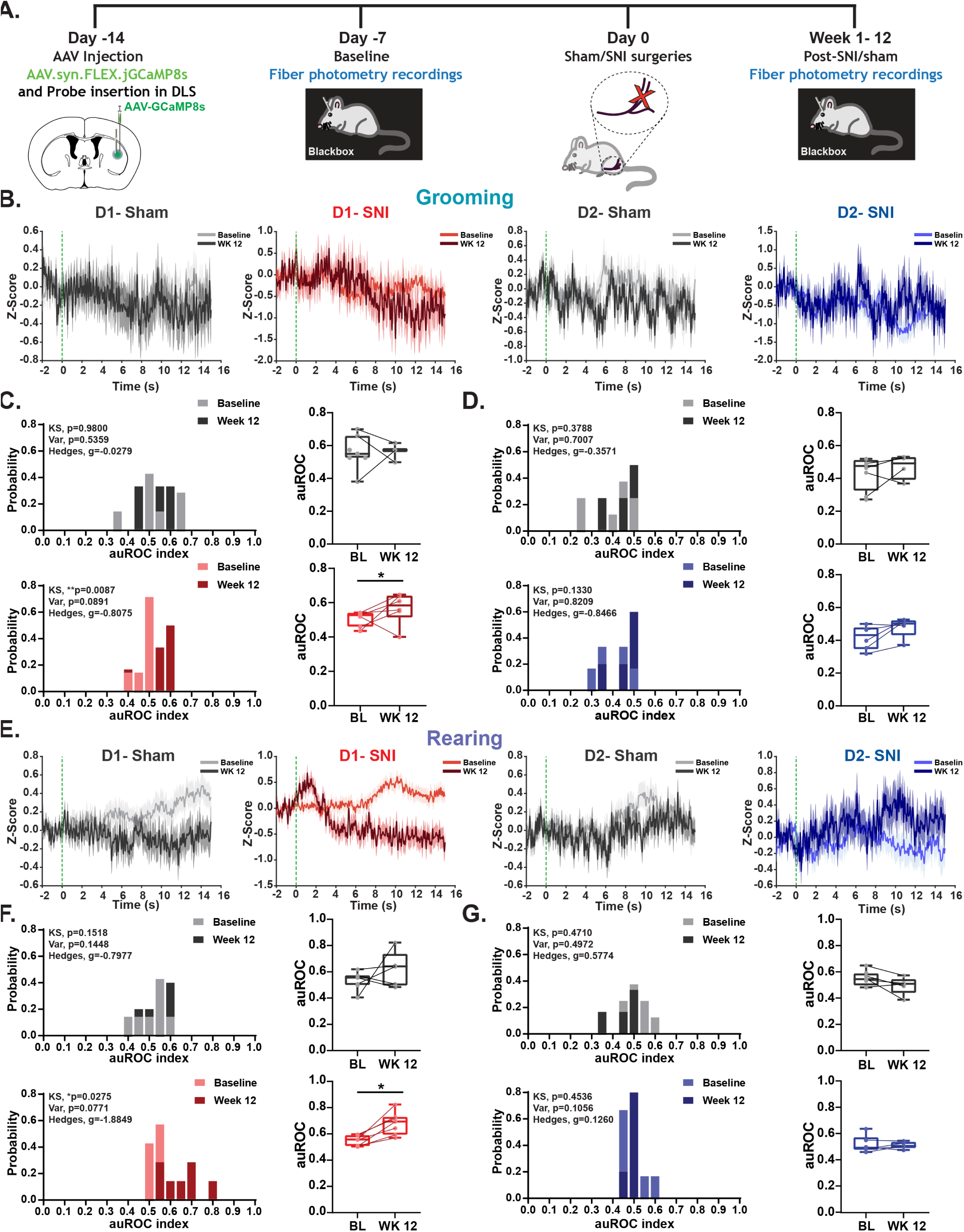
Pain-induced changes in grooming and rearing behavior modulates D1 SPN activity. **(A)** Schematic depicting timeline of AAV-GCaMP viral injection, pre-injury fiber photometry recording, and post-injury photometry recording during naturalistic behaviors in the Blackbox. **(B)** Peri-stimulus time histogram (PSTH) of mean traces sham (gray) and SNI (D1- red, D2- blue) animals comparing baseline (lighter color) to Week 12 (darker color) during grooming behavior in the Blackbox apparatus. Calcium signals are aligned to the onset of grooming (t=0). A 1–20s duration filter was applied to the PSTH. Z-score was baseline corrected. The shaded region represents SEM. **(C)** (*Left*) Probability Histograms of auROC values across baseline compared to week 12 of sham (gray) and SNI mice (red) corresponding to grooming behavior. (*Right*) auROC values of sham (gray) and SNI (red) of D1 calcium activity from baseline to Week 12. There were differences in auROC distribution in the SNI condition (D_(7,6)_ = 0.8333, **p = 0.0087; Kolmogorov-Smirnov (KS) Test). There were no differences in variance (p = 0.0891; Variance Test (Levene’s/Var-Test)). Effect size, Hedge’s g = -0.81. N(BL) = 7, N=(WK12) = 6. There were no differences in distribution of auROC values in the sham condition or in variance. Effect size, g = -0.0279. N(BL) = 7 and N(WK12) = 3. **(D)** (*Left*) Probability Histograms of auROC values across baseline compared to week 12 of sham (gray) and SNI mice (blue) corresponding to grooming behavior. (*Right*) auROC values of sham (gray) and SNI (red) of D2 calcium activity from baseline to Week 12. There were no differences in the sham or SNI condition. Sham: effect size, g = -0.36; N(BL) = 8 and N(WK12) = 4. SNI: effect size, g = -0.85; N(BL) = 6 and N(WK12) = 5. **(E)** PSTH of mean traces sham (gray) and SNI (D1- red, D2- blue) animals comparing baseline (lighter color) to Week 12 (darker color) during rearing behavior in the Blackbox apparatus. Calcium signals are aligned to the onset of rearing (t=0). A 1–20s duration filter was applied to the PSTH. Z-score was baseline corrected. The shaded region represents SEM. **(F)** (*Left*) Probability Histograms of auROC values across baseline compared to week 12 of sham (gray) and SNI mice (red) corresponding to rearing behavior. (*Right*) auROC values of sham (gray) and SNI (red) of D1 calcium activity from baseline to Week 12. There were differences in auROC distribution in the SNI condition (D_(7,7)_ = 0.7143, *p = 0.0275; KS Test). There were no differences in variance (p = 0.0771; Var-Test). Effect size, g = -1.88. N(BL) = 7, N(WK12) = 7. There were no differences in distribution of auROC values in the sham condition or in variance. Effect size, g = -0.80. N(BL) = 7 and N(WK12) = 5. **(G)** (*Left*) Probability Histograms of auROC values across baseline compared to week 12 of sham (gray) and SNI mice (blue) corresponding to rearing behavior. (*Right*) auROC values of sham (gray) and SNI (red) of D2 calcium activity from baseline to Week 12. There were no differences in the sham or SNI condition. Sham: effect size, g = 0.5774; N(BL) = 8 and N(WK12) = 6. SNI: effect size, g = 0.13; N(BL) = 6 and N(WK12) = 5.

### Acute stages of neuropathic pain confer plasticity changes in corticostriatal pathways

Thus far, our results demonstrate that neuropathic pain alters sensory-evoked response, paw and limb dynamics, and complex, naturalistic behaviors that are all correlated with modulations in D1 and D2 population activity throughout acute and chronic pain stages. We then sought to identify the synaptic mechanisms that may drive this shift. Our initial anatomical (**Supplementary Fig. 1D**) and cFos experiments (**Fig.1A**) showed that evoked pain increases neural activation in both the DLS and S1HL. Consequently, we targeted this specific S1HL-DLS corticostriatal pathway to determine if neuropathic pain induces cell-type specific plasticity changes at these synapses. We stereotaxically injected channelrhodopsin-2 (pAAV-ChR2-EYFP) unilaterally into the right hemisphere of the S1HL of D1-Cre::Ai14 and A2a-Cre::Ai14 mice, allowing expression into the corticostriatal terminals (**Fig. 4A**). In acute *ex vivo* slices, tdtomato-labeled D1- or D2-SPNs in the anterior DLS (AP from bregma: 0.5 to -0.7mm) were targeted. Whole cell current-clamp recordings were performed without inhibitory blockers to mimic natural physiological responses to optogenetic stimulation of axon terminals from cortical inputs into the DLS^49^. We employed a set of protocols to patched cells: single pulse, paired-pulse, and train stimulation to measure synaptic responses with optogenetic stimulation. To confirm our electrophysiological results were driven by biological plasticity rather than viral expression variability, we validated ChR2 expression via immunostaining (**Supplementary Fig. 5C**). Correlational analysis revealed no relationship between postsynaptic (PSP) amplitude and corrected total cell fluorescence specific to the recording area of the DLS area (**Supplementary Fig. 5D**), confirming our functional findings are independent of ChR2 expression levels. We first evaluated if there were differences in naïve controls and found no differences in intrinsic properties of rheobase, after-hyperpolarization (AHP), or resting membrane potential (RMP) (**Supplementary Fig.4A**). However, in response to single pulse, D1-SPNs had a higher PSP amplitude compared to D2-SPNs (**Supplementary Fig. 4B**). After employing paired-pulse and train stimulation protocols, there were no differences between D1- and D2-SPNs (**Supplementary Fig.4C-D**) indicating there are no plasticity changes in naïve conditions between S1-D1 and S1-D2 pathways.

**Figure 4:**
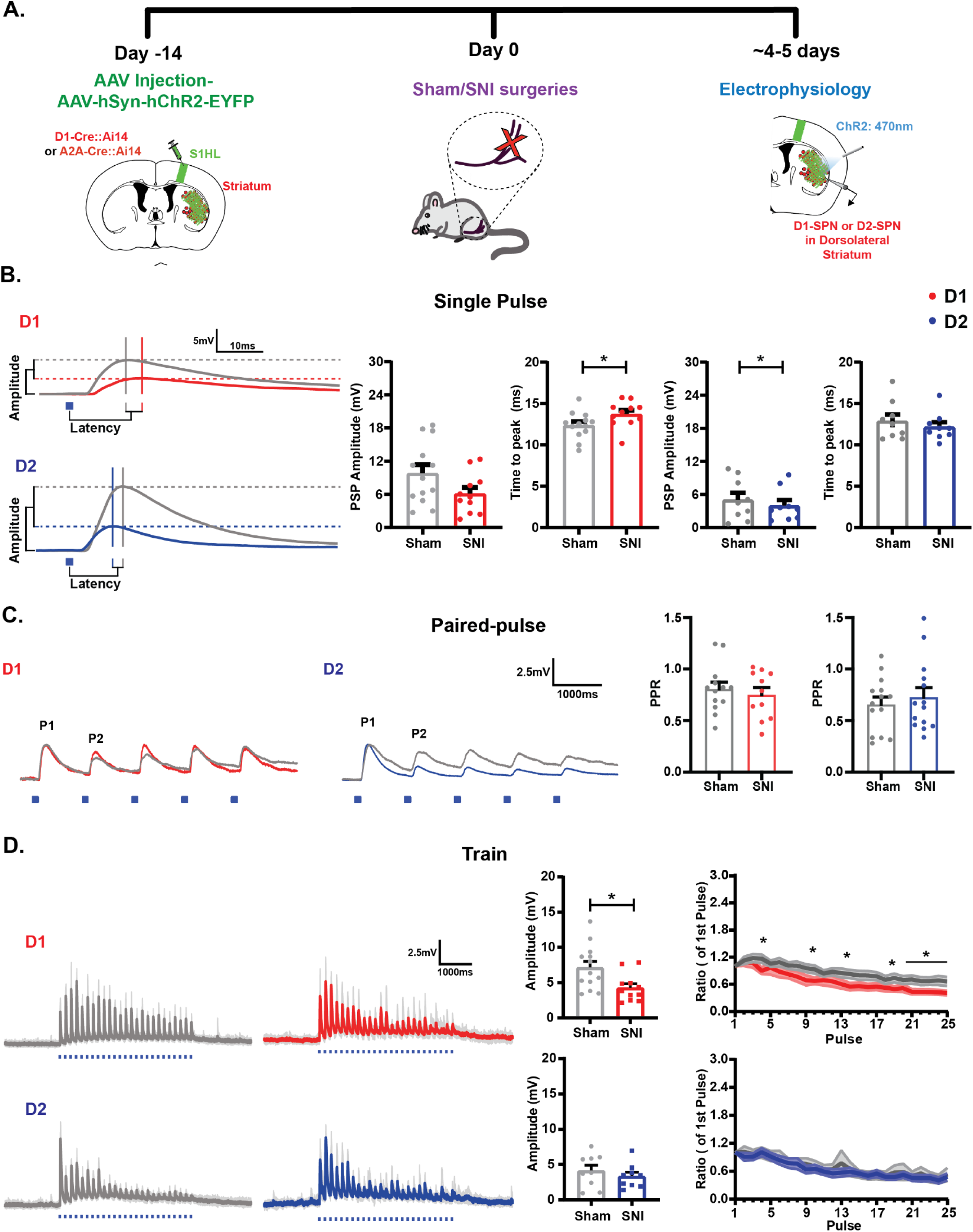
Neuropathic pain induces plasticity changes in the S1-D1 pathway during the acute phase. **(A)** Schematic depicting timeline of optogenetic electrophysiological experiments during acute phases of pain. **(B)** (*Left*) Representative schematic of S1HL-D1 (red) and S1HL-D2(blue) traces from sham and SNI mice resulting from single pulse protocol. (*Right*) Quantification of amplitude of postsynaptic (PSP) and time to peak at 5 days post-injury. There were no differences in D1 PSP amplitude. There were significant differences between D1 sham and SNI in time to peak (t_(22)_ = 2.094, p = 0.0480). There were significant differences between D2 sham and SNI conditions in PSP amplitude (t_(26)_ = 2.435, *p = 0.0221). There were no differences in time to peak. Unpaired t-test. N = 4-5 animals for sham and SNI. Error bars represent +/- SEM. **(C)** (*Left*) Representative schematic of S1HL-D1 (red) and S1HL-D2(blue) traces from sham and SNI mice resulting from paired-pulse protocol. (*Right*) Quantification of paired-pulse ratio at 5 days post-injury. There were no differences in D1 PSP amplitude. There were no differences in PPR in either D1 or D2 pathways. Unpaired t-test. N = 4-5 animals for sham and SNI. Error bars represent +/- SEM. **(D)** (*Left*) Representative schematic of S1HL-D1 (red) and S1HL-D2(blue) traces from sham and SNI mice resulting from train protocol. (*Right*) Quantification of amplitude of postsynaptic (PSP) of averaging all 25 pulses and ratio at 5 days post-injury. There were significant differences in D1 train PSP (t_(22)_ = 2.613, p = 0.0159; Unpaired t-test). There were significant differences between D1 sham and SNI in ratio of train pulses with an interaction effect (F_ConditionxPulse(24,528))_ = 1.625, *p = 0.0316) and main effect of pulse (F_Pulse(4.811,105.8)_ = 42.13, ****p< 0.0001). Two-way ANOVA with Sidak multiple comparisons. There were no significant differences in the D2 pathway. N = 4-5 animals for sham and SNI. Error bars represent +/- SEM.

Our previous data revealed that stimulus-evoked pain behavior resulted in modulation of D1- and D2-SPN populations during week 1 post-SNI (**Supplementary Fig. 1B**). Therefore, we evaluated these cell-types during the acute phase of pain (5-7 days). We analyzed the intrinsic properties of both cell types by hyperpolarizing and depolarizing current steps (**Supplementary Fig. 4E**) and found no differences between sham and SNI in either D1- or D2-SPNs (**Supplementary Fig. 4F**). In response to single pulse, D1-SPNs had no differences in PSP amplitude, but had significantly longer time to peak in the pain condition (**Fig. 4B**). However, D2-SPNs in the SNI condition had significantly lower PSP amplitude compared to sham controls without any differences in time to peak. In response to paired-pulse, there were no differences between sham and SNI conditions in either D1- or D2-SPNs (**Fig. 4C**). During 25 Hz train stimulation, however, D1-SPNs showed a significant decrease in PSP amplitude compared to shams (**Fig. 4D**). Furthermore, there was an interaction in ratio of train stimulation pulses (*p = 0.0316) and alongside a main effect of pulse (****p<0.0001), indicating diverging synaptic dynamics. Post-hoc analysis revealed SNI had significantly reduced PSP ratio at several intermittent time points with deficits becoming highly consistent during the late phase of the high-frequency train. Conversely, there were also no differences in D2-SPNs during train stimulation. Taken together, SNI alters short-term plasticity in the S1-D1 pathway and prevents the sustainability of normal synaptic drive while the S1-D2 pathway drives the postsynaptic response without any changes in synaptic plasticity during this stage.

### Transition stages of pain invoke depressing synapses in the S1-D2 pathway

To examine if the alterations in the S1-D1 and S1-D2 pathway are persistent during chronic phases of pain, we examined postsynaptic responses and plasticity changes during the transition phase of 25 days and late chronic time point of 12 weeks post-SNI (**Fig. 5A**). We applied the same protocols from the previous experiment. Similar to the acute stage of pain, during 25-days post-SNI, there were no differences in intrinsic properties in either D1- or D2-SPNs (**Supplementary Fig.5A**). However, unlike the acute stages, D1-SPNs did not show any differences in PSP amplitude or time to peak in response to single pulse (**Fig. 5B**). In contrast, SNI D2-SPNs had significantly higher PSP amplitude compared to sham controls, opposite of the acute stage. During paired-pulse protocol, the S1-D1 pathways remained unaltered, but the S1-D2 pathway in the pain condition exhibited a lower paired pulse ratio compared to shams (**Fig. 5C**). Similarly, train stimulation produced no alterations in the S1-D1 pathway whereas the S1-D2 pathway exhibited higher overall PSP amplitude and a lower train ratio, complete with an interaction effect (***p = 0.0008) and main effect of condition (*p = 0.0463) (**Fig. 5D**). These results illustrate that neuropathic pain elicits depressing synapses specifically in the S1-D2 pathway during the transition stages of pain. To provide a full picture of the late chronic stages of pain and to coincide with our fiber photometry results, we also investigated physiological changes in the S1-D1 and S1-D2 pathway at 12 weeks post-SNI. We analyzed the intrinsic properties of both S1-D1 and S1-D2 pathways and found no differences between sham and SNI conditions (**Supplementary Fig. 5B**). Contrary to both the acute phase and transition phase, there were no differences seen in either the S1-D1 or S1-D2 pathway in either post-synaptic response or plasticity changes at 12-weeks post-SNI (**Fig. 5E-G**) indicating that this circuit is critical for the transition from acute to chronic pain, rather than its long-term maintenance.

**Figure 5:**
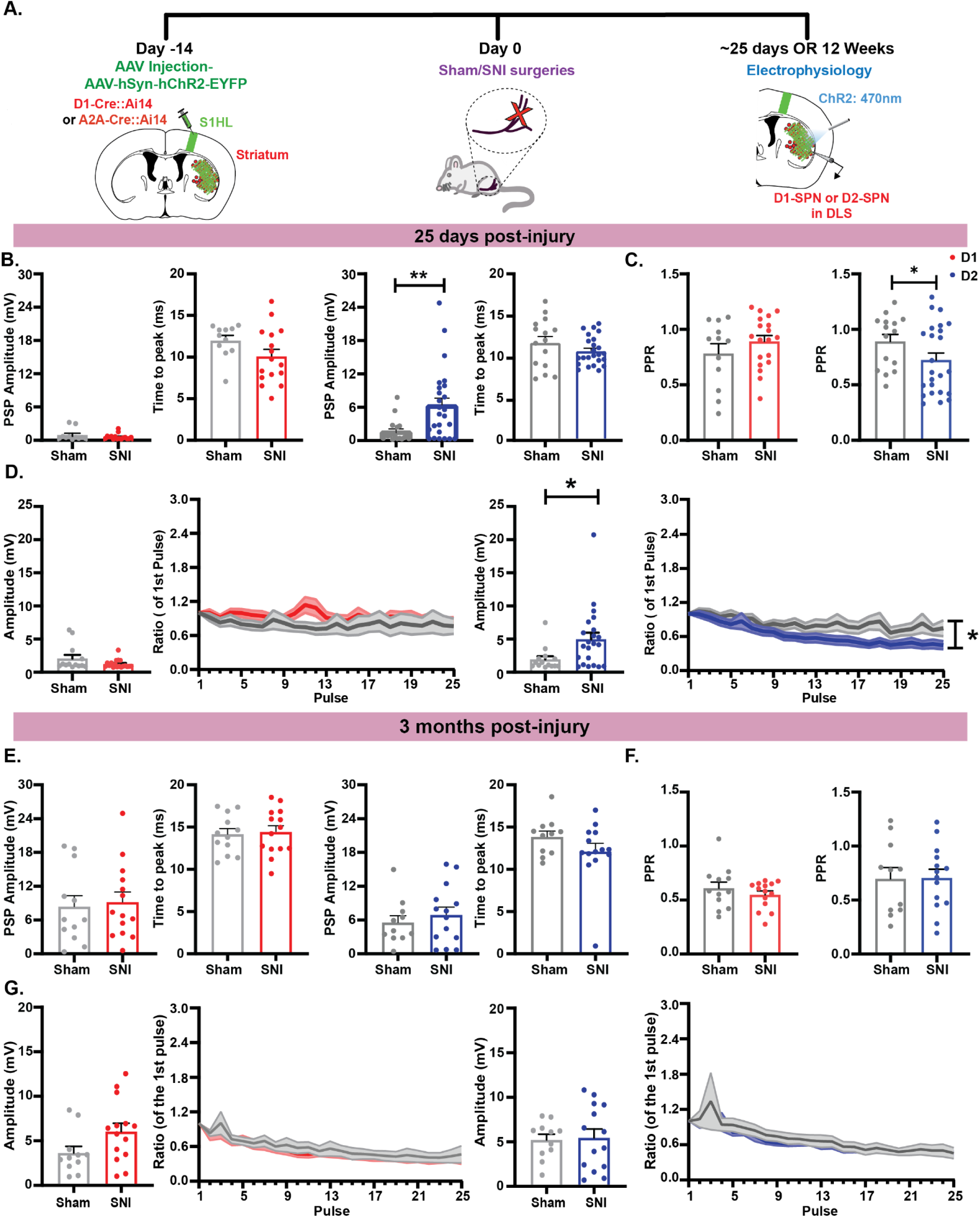
Plasticity changes occur in the S1-D2 pathway during transition pain phase. **(A)** Schematic depicting timeline of optogenetic electrophysiological experiments during transition and chronic phases of pain. **(B)** Quantification of amplitude of postsynaptic (PSP) and time to peak at 25 days post-injury during single pulse protocol. There were no differences in S1-D1 PSP amplitude or time to peak. There were significant differences between S1-D2 sham and SNI in PSP amplitude (t_(41)_ = 3.039, *p = 0.0041). There were no differences in S1-D2 time to peak. Unpaired t-test. N = 5-6 animals for sham and SNI. Error bars represent +/- SEM. **(C)** Quantification of paired-pulse ratio 25 days post-injury. There were no differences in D1 PPR. There were significant differences in S1-D2 PPR (t_(37)_ = 1.782, *p = 0.0415). Unpaired t-test. N = 5-6 animals for sham and SNI. Error bars represent +/- SEM. **(D)** Quantification of amplitude of postsynaptic (PSP) of averaging all 25 pulses and ratio at 25 days post-injury during train protocol. There were no significant differences in S1-D1. There were significant differences in S1-D2 in PSP amplitude (t_(34)_ = 2.391, *p = 0.0225; Unpaired t-test). There were also significant differences in S1-D2 train ratio with an interaction effect (F_ConditionxPulse(24,864)_ = 2.202, ***p = 0.0008), main effect of pulse (F_Pulse(24,864)_ = 16.16, ****p< 0.0001), and main effect of condition (F_Condition(1,36)_ = 0.0463, *p = 0.0463). Two-way ANOVA with Sidak multiple comparisons. N = 5-6 animals for sham and SNI. Error bars represent +/- SEM. **(E)** Quantification of amplitude of postsynaptic (PSP) and time to peak at 12 weeks post-injury during single pulse protocol. There were no differences in S1-D1 or S1-D2 pathways. N = 4-5 animals for sham and SNI. Error bars represent +/- SEM. **(F)** Quantification of paired-pulse ratio 12 weeks post-injury. There were no differences in S1-D1 or S1-D2 pathways. N = 4-5 animals for sham and SNI. Error bars represent +/- SEM. **(G)** Quantification of amplitude of postsynaptic (PSP) of averaging all 25 pulses and ratio at 12 weeks post-injury during train protocol. There were no differences in S1-D1 or S1-D2 pathways. N = 4-5 animals for sham and SNI. Error bars represent +/- SEM.

### Neuropathic pain promotes unbiased macroscale diversity in movement

Because our physiological data revealed profound synaptic divergence specifically during the acute and transition phases of pain, we sought to determine if this circuit instability drives hidden, macroscale movement patterns missed by supervised learning. Therefore, we took an unsupervised learning approach to our behavioral studies and applied Variational Embeddings of Animal Motion (VAME) to the same Blackbox data set which allows for probabilistic deep learning of behavioral states using pose estimation tools^48^. The model identified 15 behavioral motifs clustered into five communities: C1, low-speed locomotion; C2, quiescence/resting; C3, micromovements/fine-motor investigation; C4, exploratory movement (sniffing/rearing); and C5, high-speed locomotion. Evaluating time spent in each community revealed that SNI mice exhibited fewer bouts in C2 (interaction *p = 0.0488, main effect *p = 0.0269) and C4 (main effect **p = 0.0017) relative to shams, reflecting reduced resting and exploratory behavior (**Supplementary Fig. 6A**). Communities 3 and 5 showed main effects of time (*p = 0.0470 and *p = 0.0411, respectively), indicating a longitudinal decrease across both groups. To assess shifting behavioral strategies, we constructed a grammar-based map using 5×5 transition matrices reduced via UMAP (**Fig. 6A**). Permutation testing revealed that SNI mice exhibited significant behavioral divergence specifically at Week 3 (**Fig. 6B**), accompanied by higher behavioral entropy. There was no divergence that occurred at Weeks 1 or 12 (**Supplementary Fig. 6B**). Granular analysis of Week 3 transition probabilities (**Fig. 6C)** demonstrated that SNI animals had increased transitions from C2→C1, C3→C4, C3→C5, and C5→C4, alongside decreased transitions from C1→C2, C5→C1, C4→C3, and C5→C3 (with distinct shifts at Weeks 1 and 12; **Supplementary Fig. 6C**). Collectively, this unsupervised approach reveals that the transition phase of chronic pain (Week 3) is marked by maximal behavioral volatility and complex, shifting movement strategies, mirroring the synaptic instability observed in the corticostriatal circuit at this time point.

**Figure 6:**
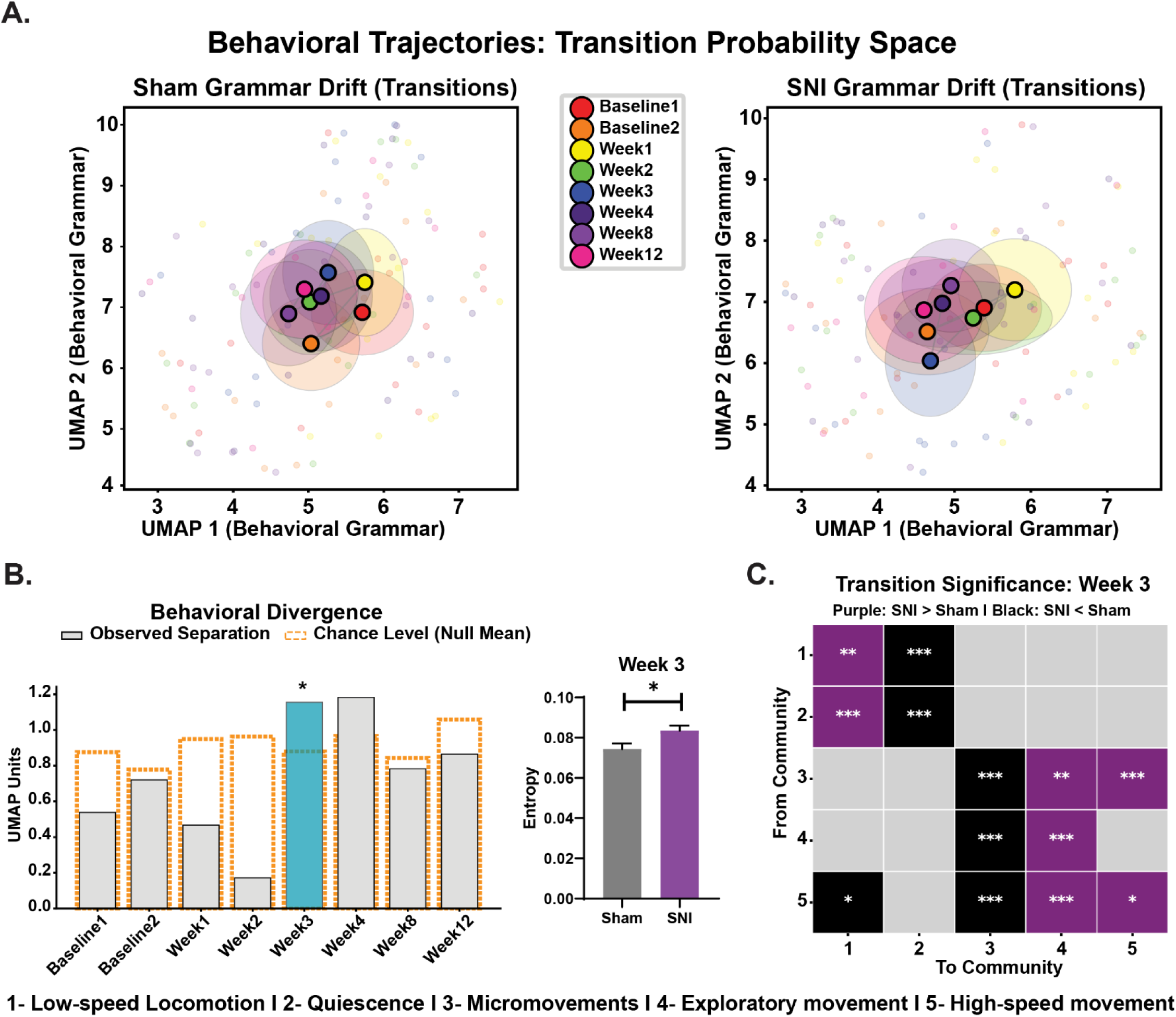
Neuropathic pain reveals transition phase with the most behavioral divergence. **(A)** Longitudinal behavioral grammar trajectories in transition probability manifold. Two-dimensional Uniform Manifold Approximation and Projection (UMAP) embedding of the 25-dimensional Markovian transition probability matrices (5×5 community transitions) across 218 sessions in Sham (left) and SNI (right) mice. Small dots denote individual recording sessions across longitudinal timepoints (Baseline 1 through Week 12; colored according to timeline key). Large, outlined circles represent group centroids at each timepoint, dashed ellipses indicate +/-2 SEM along each manifold dimension, and dashed gray lines trace the mean longitudinal trajectory across weeks. Note the divergence of the SNI trajectory away from the stable Sham manifold baseline, particularly around Week 3 (blue). **(B)** Quantitative behavioral divergence and entropy collapse at Week 3. (*Left*) Group-level behavioral grammar divergence between Sham and SNI cohorts across timepoints, quantified as the Euclidean distance between cohort centroids in UMAP space. Solid gray bars indicate the observed Euclidean separation; dashed open bars indicate the chance level derived from the mean of the null distribution generated by label permutation (N = 1,000 iterations). Week 3 (blue bar) exhibited significant divergence from healthy controls (*p<0.05, non-parametric permutation test). (Right) Behavioral sequencing entropy at Week 3 post-injury. Bars/points represent mean row-wise Shannon entropy computed from individual 5×5 transition probability matrices for Sham and SNI mice. SNI animals demonstrated altered behavioral entropy compared to Sham controls (t_(26)_ = 2.413, *p = 0.0232; Unpaired t-test), indicating a shift toward restricted, stereotyped transition syntax. N = 15 for sham and N = 13 for SNI. Error bars represent +/- SEM. **(C)** Reorganized community transition architecture at Week 3. Heatmap depicting pairwise transition significance between the five behavioral communities (C1: Low-speed Locomotion; C2: Quiescence; C3: Micromovements; C4: Exploratory Movement; C5: High-speed Locomotion) at Week 3 post-injury. Rows indicate the originating community (“From”), and columns indicate the terminating community (“To”). Colors indicate the direction of significant probability alterations in SNI relative to Sham: purple indicates significant increase in transition frequency (SNI>Sham), black indicates a significant reduction (SNI<Sham), and gray denotes non-significant transitions (NS). Significance was determined via two-tailed Pearson’s *X^2^* contingency tests with Laplace (+1) continuity smooth (*p<0.05, **p<0.01, ***p<0.001).

### Unbiased movement strategies are attuned to D1 and D2 SPN activity

Having established Week 3 as a critical transition phase characterized by peak behavioral divergence and entropy, we sought to determine how SPN activity maps onto these shifting movement strategies. Building on our supervised learning photometry results (**Fig. 3**), we correlated D1- and D2-SPN calcium dynamics with each VAME-identified community at Week 3 (**Fig. 7A,D**). We first focused on Community 3 and 5 because these two communities were the most involved in week 3 transition probabilities. We applied a similar auROC analysis as we did in Figure 3 and found that within Community 3, D1-SPNs remain stable in both the pain condition and the sham controls (**Fig. 7B**). Conversely, we found that the D2-SPNs in the pain condition transitioned from a more suppressed state to a more active state, becoming strong encoders of micromovements at week 3 (KS, *p = 0.0122; *g* = -1.716) compared to sham controls (**Fig. 7C**). Within Community 5, the D1-SPN population exhibited similar distribution changes across time in both sham and SNI mice, yielding moderate effect sizes without any statistical significance (**Fig. 7E**). However, the D1-SPN population did have significant variance (*p = 0.0166) pointing to the SNI group becoming more scattered and implying that chronic pain introduces heterogeneity in the D1 population during high-speed locomotion. The D2 population also did not have significant differences in their distributions between baseline and week 3 (**Fig. 7F**), but SNI animals did have a large effect (*g* = 0.9664) suggesting that D2 neurons are losing their tuning to high-speed locomotion. We also investigated the other communities and found that there were no chronic pain differences in Community 2 (**Supplementary Fig. 7A-C**). In Community 4, D1-SPNs in the pain condition were found to significantly shift to become more suppressed (KS, *p = 0.0275; *g* = 1.0183) indicating D1 neurons do not sustain encoding for exploratory behaviors (**Supplementary Fig. 7E**) while D2-SPNs remain stable (**Supplementary Fig. 7F**). For community 1, D1-SPNs do not show significant shifts over time in either sham controls or SNI animals; however, both groups show significant variance (**p = 0.0094 and *p = 0.0111, respectively) (**Supplementary Fig. 6E**). D2-SPNs demonstrate functional stability in the sham controls while the pain condition revealed trending decreases in encoding for community 1 (KS, p = 0.0766, g = 1.0362) and significant variability (*p = 0.0282) also pointing to heterogeneity in the D2 population during low-speed walking (**Supplementary Fig. 6F**). Collectively, this unbiased approach demonstrates that the peak behavioral entropy observed at Week 3 is underpinned by a pathway reorganization, where D2-SPNs become heavily tuned to micromovements while D1-SPNs simultaneously uncouple from exploratory behaviors. Ultimately, these findings emphasize that the complex movement shifts induced by neuropathic pain are driven by distinctly different adaptations in D1- and D2-SPNs, with each population independently reorganizing around different macroscale behavioral states.

**Figure 7:**
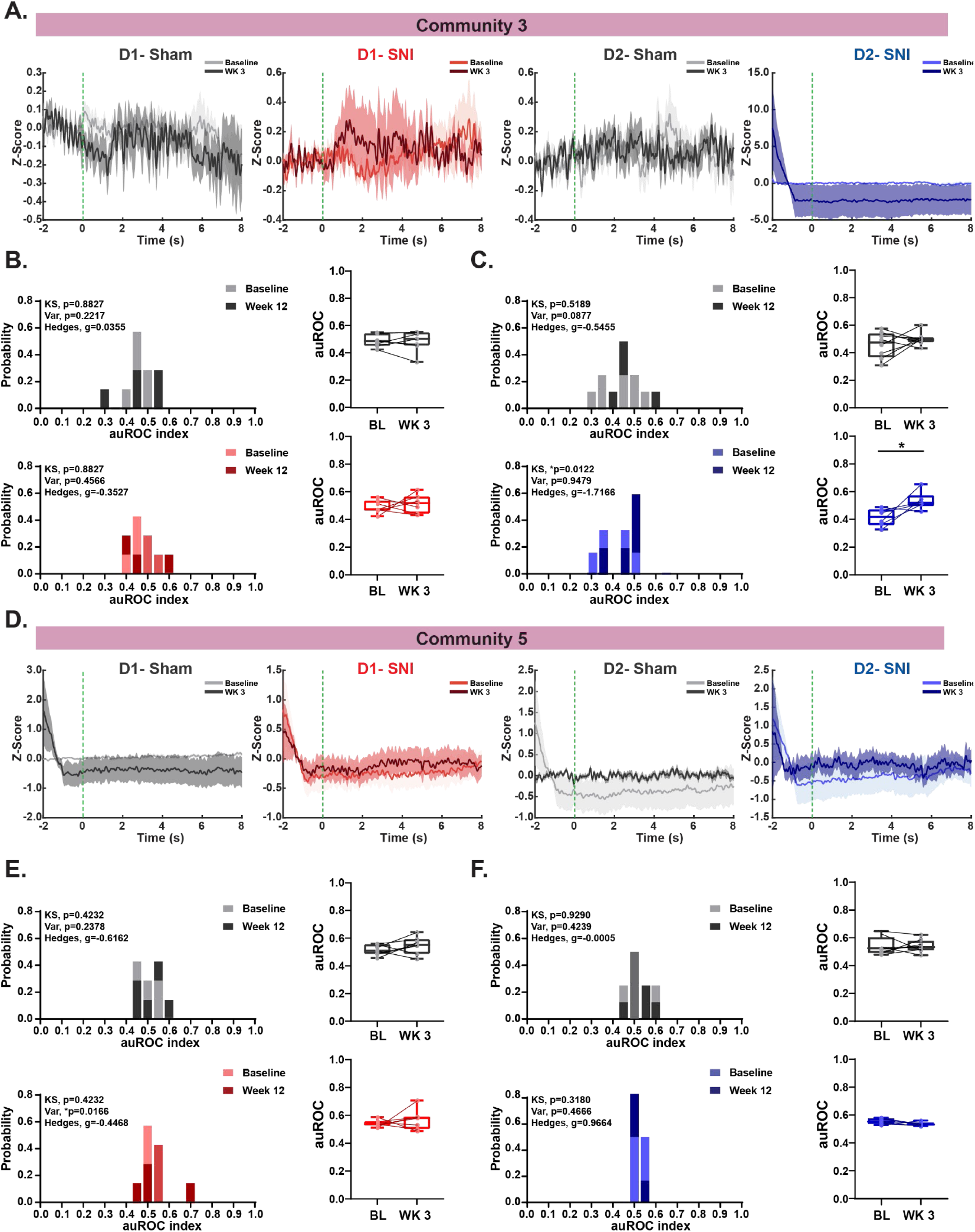
Micromovements are modulated by D2-SPNs. **(A)** Peri-stimulus time histogram (PSTH) of mean traces sham (gray) and SNI (D1- red, D2- blue) animals comparing baseline (lighter color) to Week 3 (darker color) during community 3. Shaded region represents SEM. **(B)** (*Left*) Probability Histograms of auROC values across baseline compared to week 3 of sham (gray) and SNI mice (red) corresponding to community 3. (*Right*) auROC values of sham (gray) and SNI (red) of D1 calcium activity from baseline to Week 3. There were no differences in auROC distribution in either sham or SNI. Sham: effect size, g = 0.0355, N(BL) = 7, N(WK3) = 7. SNI: effect size, g =-0.3527, N(BL) = 7 and N(WK3) = 7. **(C)** (*Left*) Probability Histograms of auROC values across baseline compared to week 3 of sham (gray) and SNI mice (blue) corresponding to community 3. (*Right*) auROC values of sham (gray) and SNI (red) of D2 calcium activity from baseline to Week 3. There were differences in auROC distribution in the SNI condition (D_(6,6)_ = 0.8333, *p =0.0122; KS Test). There were no differences in variance (p = 0.9479). Effect size, Hedge’s g = -1.7166. N(BL) = 6, N(WK3) = 6. There were no differences in distribution of auROC values in the sham condition or in variance. Effect size, g = -0.5455. N(BL) = 8 and N(WK3) = 8. **(D)** Peri-stimulus time histogram (PSTH) of mean traces sham (gray) and SNI (D1- red, D2- blue) animals comparing baseline (lighter color) to Week 3 (darker color) during community 5. Shaded region represents SEM. **(E)** (*Left*) Probability Histograms of auROC values across baseline compared to week 3 of sham (gray) and SNI mice (red) corresponding to community 5. (*Right*) auROC values of sham (gray) and SNI (red) of D1 calcium activity from baseline to Week 3. There were no differences in auROC distribution in either sham or SNI. There was significant variance (F_(6,6)_ = 0.1097, *p =0.0166; Var Test). Effect size, g = -0.4468, N(BL) = 7, N(WK3) = 7. There were no differences in the sham condition. Effect size, g =-0.6162, N(BL) = 7 and N(WK3) = 7. **(F)** (*Left*) Probability Histograms of auROC values across baseline compared to week 3 of sham (gray) and SNI mice (red) corresponding to community 5. (*Right*) auROC values of sham (gray) and SNI (blue) of D2 calcium activity from baseline to Week 3. There were no differences in auROC distribution in either sham or SNI. Sham: effect size, g = -0.0005. N(BL) = 8, N(WK3) = 8. SNI: effect size, g = 0.9664. N(BL) = 6 and N(WK3) = 6.

## Discussion

The goal of this study was to investigate the longitudinal progression of pain signatures by uncovering the striatal circuit dynamics through a sensorimotor lens. Up until now, most preclinical studies have focused on the ventral striatum to investigate distinct changes in threshold, aversion, and motivational effort^57, 58^, often limiting chronic pain models to a maximum of 3 weeks post-injury. Despite overwhelming clinical evidence demonstrating the Dstr at the center of the pain circuit^13, 15, 59–61^, this sensorimotor hub has never been tested within this framework or along this time progression. Our work functionally defined the temporal evolution of neuropathic pain into distinct phases of acute (Week 1), transition (Week 3), and late chronic (Week 12). While most preclinical studies define chronic pain at 2-3 weeks post-injury, we argue that clinically defined chronic pain is closer to the 3-month (12 week) timepoint, and that 3 weeks post-injury represents a volatile transition phase. Our multi-perspective behavioral analysis exposed how different types of behavior adapt along this pain time course. Evoked reflexive behaviors were characterized by persistent mechanical hypersensitivity that was uncoupled from withdrawal vigor during the entire progression. In expanding the behavioral lens to investigate complex naturalistic behaviors, we identified maladaptive reductions in self-soothing and exploratory behaviors that emerged strongly at week 12. Concurrently, our unbiased approach revealed that neuropathic pain drives distinct macroscale movement strategies with the peak of behavioral diversity occurring at Week 3. Importantly, these behavioral dynamics were reflected through changes in D1- and D2-SPN dynamics and synaptic physiology, with the height of corticostriatal plasticity changes occurring during the transition phase of week 3.

### Resolving SPN population activity vs. cortical inputs

D1- and D2-SPNs represent the majority of the striatal network (∼95%) and have distinct functional roles, suggesting that chronic pain might differentially affect these two populations to shift the balance toward avoidance behaviors that characterize chronic pain states. Indeed, we found that throughout the continuum of pain progression, the SPN population and corticostriatal pathways exhibit distinct changes in activation and plasticity. During the acute phase (week 1), both D1- and D2-SPNs demonstrate delayed temporal dynamics in response to evoked stimuli **(Supplementary Fig. 1E**) and specifically the S1HL-D1 pathway exhibits short-term depression while the S1HL-D2 pathway demonstrates reductions in postsynaptic response without any plasticity changes (**Fig. 4**). Conversely, during the transition phase (week 3), D2-SPNs take on a more active role during neuropathic pain by becoming strong encoders of micromovements (**Fig. 7**) coinciding with the S1HL-D2 pathway developing depressing synapses (**Fig. 5**). D1-SPNs lose tuning for exploratory behaviors, but unlike the D2 pathway, the S1HL-D1 pathway normalizes during this timepoint. During the late chronic phase (week 12), we find that D1-SPNs drive the chronic state by exhibiting persistent timing delays to evoked stimuli (**Fig. 1**) and becoming significantly positively modulated with grooming and rearing (**Fig. 3**). Meanwhile, the corticostriatal synaptic plasticity normalizes across both pathways suggesting that these specific synapses drive the transition to chronic pain rather than its long-term maintenance. All in all, neuropathic pain does not uniformly alter striatal output; rather, D1 and D2 SPN populations undergo divergent, functional reorganizations to encode distinct behavioral strategies.

### Striatum, pain, and mechanism for treatment

The striatum is a focal point of sensory signal and motor output, governing action selection and movement initiation. Chronic pain is not just a disorder of sensation, but of motor response which positions the DStr as an ideal candidate to mediate pain-induced changes in movement behavior, yet it remains a compelling but understudied network in chronic pain pathology. Many of the preclinical studies focused specifically on the ventral striatum demonstrating reorganization of ventral striatal pathways^62^ with changes in neurotransmitter content, and specifically implicating D2-SPNs in mediating pain hypersensitivity and depressive-like behaviors^63, 64^. The few preclinical studies that have targeted the DStr (such as orofacial pain) have also highlighted the role of D2-SPNs in inhibiting the nociceptive jaw opening reflex, implicating this indirect pathway in endogenous analgesia^28, 65^. Coinciding with this, in an acute inflammatory pain model, inhibition of D2 receptor cells in the DLS, but not D1, increases nociception, whereas activation of these cells reduces nociception^66^. Furthermore, in a neuropathic pain model (2-4 weeks post-SNI), another study substantiated that the activation of D2-SPNs weakens pain hypersensitivity^67^. Our work agrees with the previous literature regarding the active role of the DStr, particularly D2-SPNs and related corticostriatal pathways during the transition phase of 3 weeks post-SNI. However, our longitudinal work crucially reveals the active role of D1-SPNs throughout the entirety of pain progression. Because peak plasticity changes occurred in the S1HL-D1 pathway during the acute phase and D1-SPNs strongly attuned with complex behaviors in the late chronic phase, our data suggests the D1 population is a critical, previously overlooked therapeutic target for preventing pain chronification.

The divergent roles of D1 and D2 SPNs identified here also provide critical context for pharmacological pain management, particularly regarding the opioid system. The DStr is characterized by a large concentration of opiates and cannabinoids. Human imaging studies reveal high levels of pain-related dopaminergic and opioidergic expression and activation, with large overlap between these populations within the DStr, suggesting these mechanisms are involved in modulate the perception of aversive experiences related to pain^68–70^. Preclinical studies illustrate a strong functional link between D2-SPNs and the opioid system, with D2 receptors bidirectionally regulating enkephalin expression and affecting reward behavior^71^. However, other studies demonstrate a D1-pathway-specific effect associated with the acquisition of opioid-seeking behaviors, with D1 SPNs having reduced sensitivity after opioid self-administration^72^. While most of these studies were done in the ventral striatum, other research shows that within the DStr D1-pathway, repeated opioid exposure and withdrawal potentiated GABAergic striatopallidal and striatonigral transmission, which inhibited substantia nigra pars compacta dopamine release, contributing to negative affect^73^. Recent studies also highlight the opposing effects of mu-opioid receptors on D1- and D2-SPNs in opioid mediated antinociception in acute, inflammatory models in which opioid antinociception was attenuated in conditions where D1-SPNs lacked mu opioid receptors, but was enhanced when mu opioid receptors were ablated in D2-SPNs^74^. Our findings suggest that treating chronic pain effectively may require leveraging these cell-type-specific opioid mechanisms precisely when those populations are most actively driving behavioral adaptations.

### Future implications

Our work advances the field of both pain and striatal networks by unraveling how striatal circuits adapt to persistent pain through synaptic plasticity, population dynamics, and neural timing. By analyzing striatal biology at the pathway-specific and population-level, we reveal the distinct, time-dependent roles of D1 and D2 SPNs in the transition from acute to chronic pain. Furthermore, this work illuminates potential critical windows, such as the highly volatile Week 3 transition phase, laying the foundation for time-specific, circuit-precise therapeutic targets in persistent pain.

## Acknowledgements

The authors would like to thank members of the Abraira Lab: Aman Upadhyay, Joshua Thackray, and Alana Martinez for their help. This work was supported by the National Institutes of Health/Neurological Disease and Stroke (F32NS132956, A.J.G.); (R01NS124799, V.E.A.); (R01NS094450; D.J.M.); the Rita Allen Foundation and Sloan Foundation (V.E.A).

## Contributions

Conception and design were carried out by A.J.G., D.J.M., and V.E.A. Acquisition of data was performed by A.J.G. and M.G. Analysis of data was conducted by A.J.G., I.L.G., A.J.Y., X.C.Z., and A.D. The ARM equipment was built by J.B. who also provided assistance to ARM data analysis. Drafting and revision of the paper were completed by A.J.G., M.G., D.J.M., and V.E.A. Oversight was provided by D.J.M. and V.E.A.

## Corresponding authors

Correspondence to David Margolis and Victoria Abraira

## Competing interests

Provisional Patent no. 63/521,444 was filed for the ARM device on 6-16-2023. In addition, J.B. is Co-Founder of Tactorum Inc., a company that mass produces the ARM device and associated pain assessment software.

## Data availability

All data supporting the findings of this study are available within the paper and its supplementary information. All data reported in this paper will be shared by the lead contact, A.J.G. upon request.

## Code availability

All codes for supervised, unsupervised learning behavior and calcium analysis can be found at: https://github.com/margolislab/D1_D2_FP_2026

## Notes

### Competing Interest Statement

Competing interests: Provisional Patent no. 63/521,444 was filed for the ARM device on 6-16-2023. In addition, Justin Burdge is Co-Founder of Tactorum Inc., a company that mass produces the ARM device and associated pain assessment software.

### Summary of Updates

In this revision, we added new data, new analysis, updated text, added two co-authors.

