## Supplemental Figures 1-7 for "Neuropathic pain drives time-dependent reorganization of naturalistic movement and corticostriatal circuits"

### Supplementary Figure 1

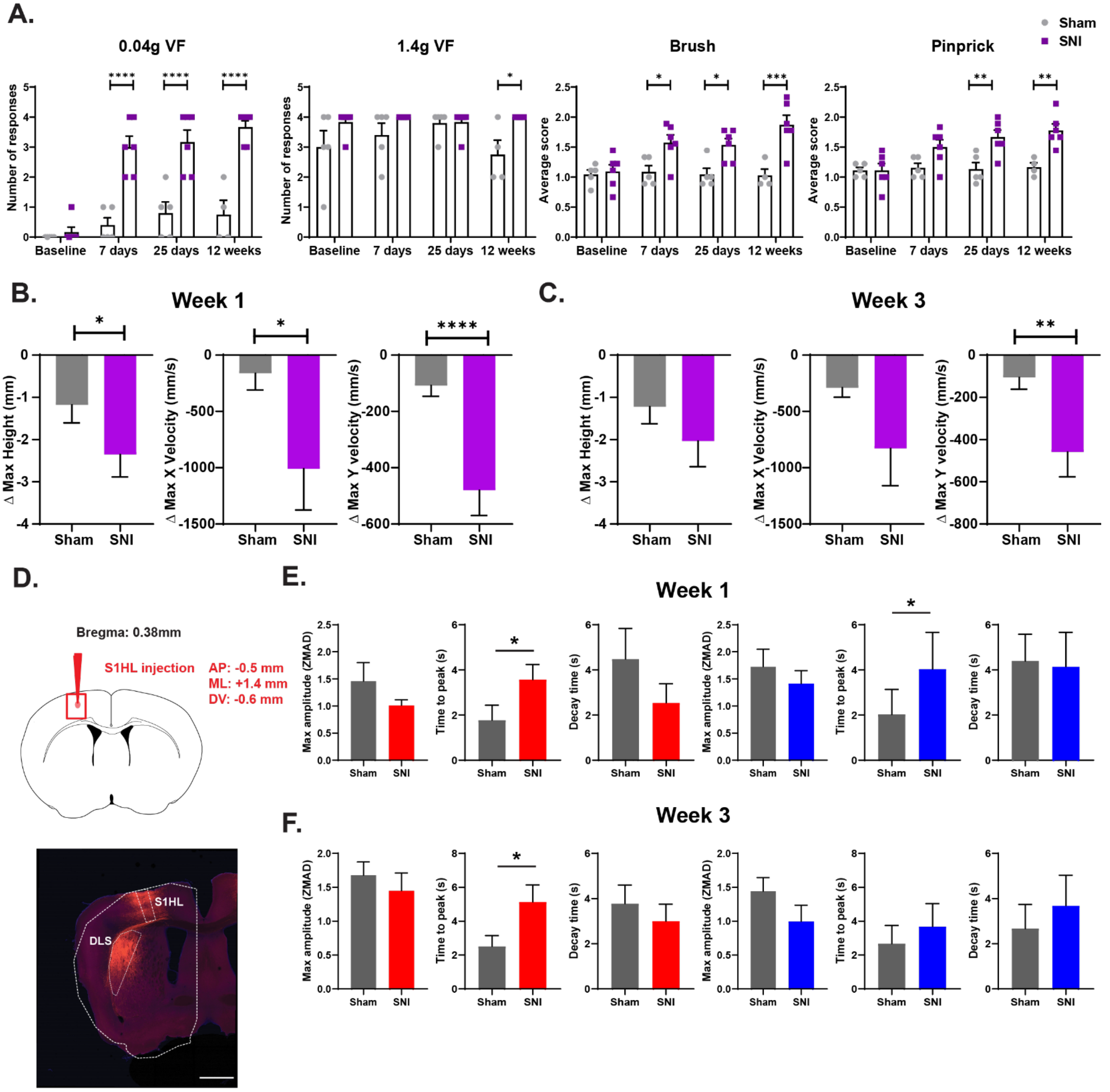

**Supplementary Figure 1**

**(A)** Manual stimulation with Von Frey (VF), brush, and pinprick to the ipsilateral injured paw during baseline, 7 days, 25 days, and 3 months post-injury. No differences were found at baseline. There were significant differences found at 0.04g VF with a significant interaction $F_{ConditionxTime(3, 26)}$ = 10.95, ****p<0.0001, significant main effect of condition $F_{condition(1, 9)}$ = 48.05, ****p<0.0001, and significant main effect of time $F_{Time(3, 26)}$ = 26.44, ****p<0.0001 and significant post-hoc comparisons at 7 days, 25 days, and 3 months (****p<0.0001). At 1.4g VF, there was a significant main effect of condition $F_{condition(1, 9)}$ = 9.660, *p = 0.0126 with significant post-hoc comparisons as 12 weeks (*p = 0.0163). For brush, there was significant interaction $F_{ConditionxTime(3, 26)}$ = 4.173, *p = 0.0154, significant main effect of condition $F_{condition(1, 9)}$ = 21.21, **p = 0.0013, and significant main effect of time $F_{Time(3, 26)}$ = 4.068, *p = 0.0170 and significant post-hoc comparisons at 7 days (*p = 0.0250), 25 days (*p = 0.0223), and 12 weeks (***p = 0.0001). For pinprick, there was significant interaction $F_{ConditionxTime(3, 26)}$ = 3.835, *p = 0.0213, significant main effect of condition $F_{condition(1, 9)}$ = 18.86, **p = 0.0019, and significant main effect of time $F_{Time(3, 26)}$ = 4.937, **p = 0.0076 and significant post-hoc comparisons at 25 days (**p = 0.0034), and 12 weeks (**p = 0.0014). Mixed-effects model (REML) with Sidak's post-hoc comparisons. N = 5 for sham and N = 6 for SNI. Error bars represent ± SEM.

**(B)** PAWS analysis of hindlimb trajectory from ARM experiments during Week 1 post-injury, when normalized to baseline (Δ = post-injury - baseline). Significant differences were observed in maximum height ( $t_{(26)}$ = 1.751, *p = 0.0459. Unpaired t-test), maximum x velocity (U = 58, *p = 0.0360), and maximum y velocity (U = 17, ****p<0.0001 Mann Whitney U). N = 15 for sham and N = 13 for SNI. Error bars represent +/- SEM.

**(C)** PAWS analysis of hindlimb trajectory from ARM experiments during Week 3 post-injury, when normalized to baseline (Δ = post-injury - baseline). No significant differences were observed in maximum height or maximum x velocity. A significant difference was found in maximum y velocity (U = 17, **p = 0.0031. Mann Whitney U). N = 11 for sham and N = 10 for SNI. Error bars represent ± SEM.

**(D)** Anterograde tracing studies using AAV-CAG-tdtomato injected into S1HL region (*top*). Image showing dense projections to the dorsolateral striatum (DLS) (*bottom*). The scale bar is 1000μm.

**(E)** Calcium activity of D1 (red) and D2 (blue) during ARM experiments at week 1 and week 3 post-injury. At week 1, there were no significant differences in maximum amplitude for either D1 or D2. There were significant differences for time to peak in D1, U = 6, *p = 0.0157; and D2, U = 10, *p = 0.0406. There are no significant differences in decay time. For week 3, there were no significant differences in maximum amplitudes for D1 or D2. There were significant differences in time to peak for D1, U = 8, *p = 0.0379, but not D2. There were no significant differences in decay time for either D1 or D2. Mann Whitney U. For D1, N = 7 for sham and N = 7 for SNI. For D2, N = 8 for sham and N = 6 for SNI. Error bars represent ± SEM.

### Supplementary Figure 2

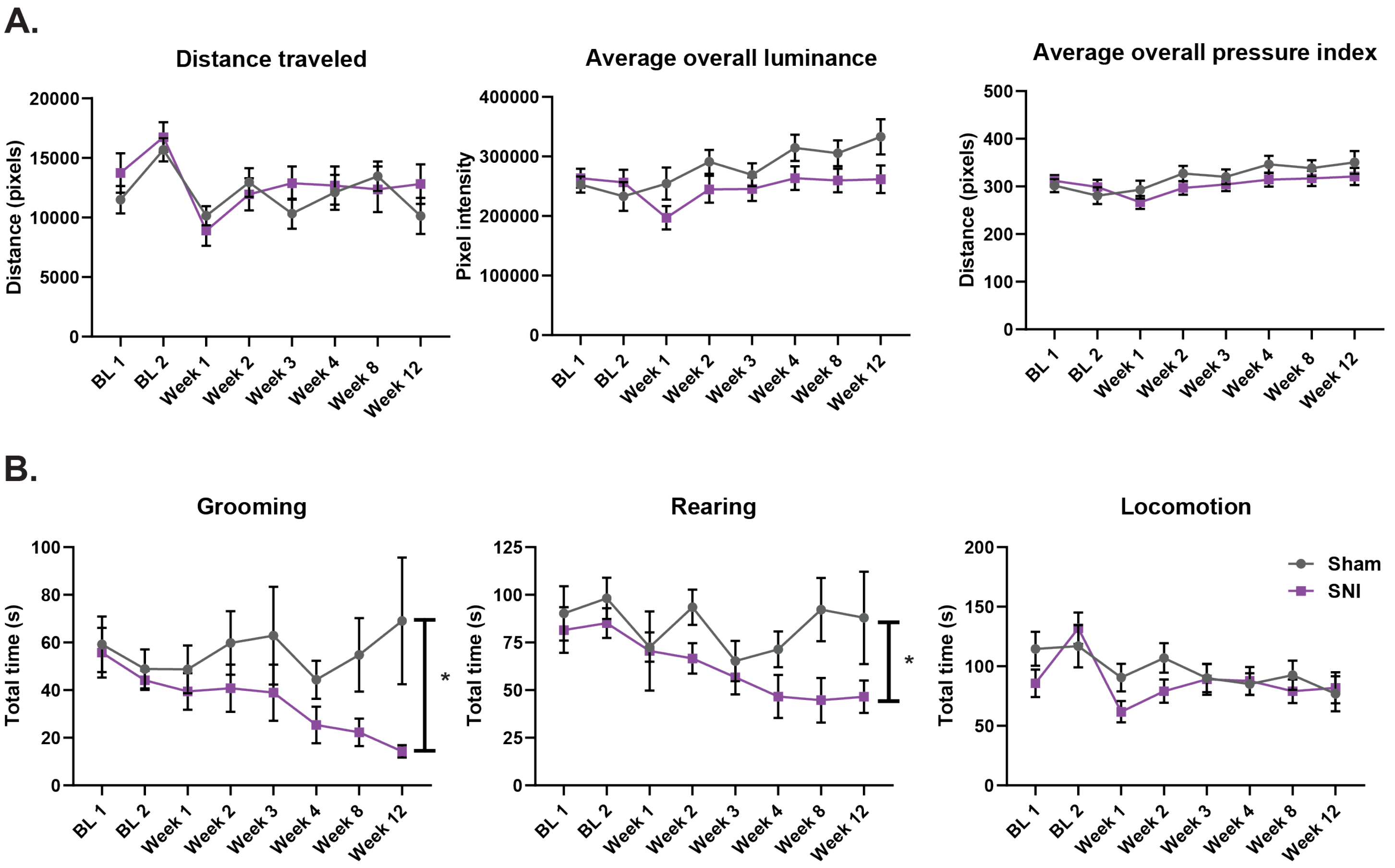

**Supplementary Figure 2**

**(A)** Parameters for overall paw dynamics in the Blackbox. There were no significant differences in distance traveled, luminance, or pressure over the course of 12 weeks. N = 12-16 for sham and N = 12-13 for SNI. Error bars represent ± SEM.

**(B)** Total time spent grooming, rearing, or locomotion extracted from supervised learning of SimBA analysis from Blackbox experiments over the course of 12 weeks. For grooming, there were significant differences with a main effect of condition, $F_{condition(1, 26)} = 5.571$, *p = 0.0260 with significant post-hoc comparisons as 12 weeks (*p = 0.0101). For rearing, there were significant differences with a main effect of condition, $F_{condition(1, 26)} = 5.842$, *p = 0.0230, and a main effect of time, $F_{Time(1, 26)} = 2.090$, *p = 0.0468. There were no significant differences in locomotion. N = 12-16 for sham and N = 12-13 for SNI. Error bars represent ± SEM.

### Supplementary Figure 3

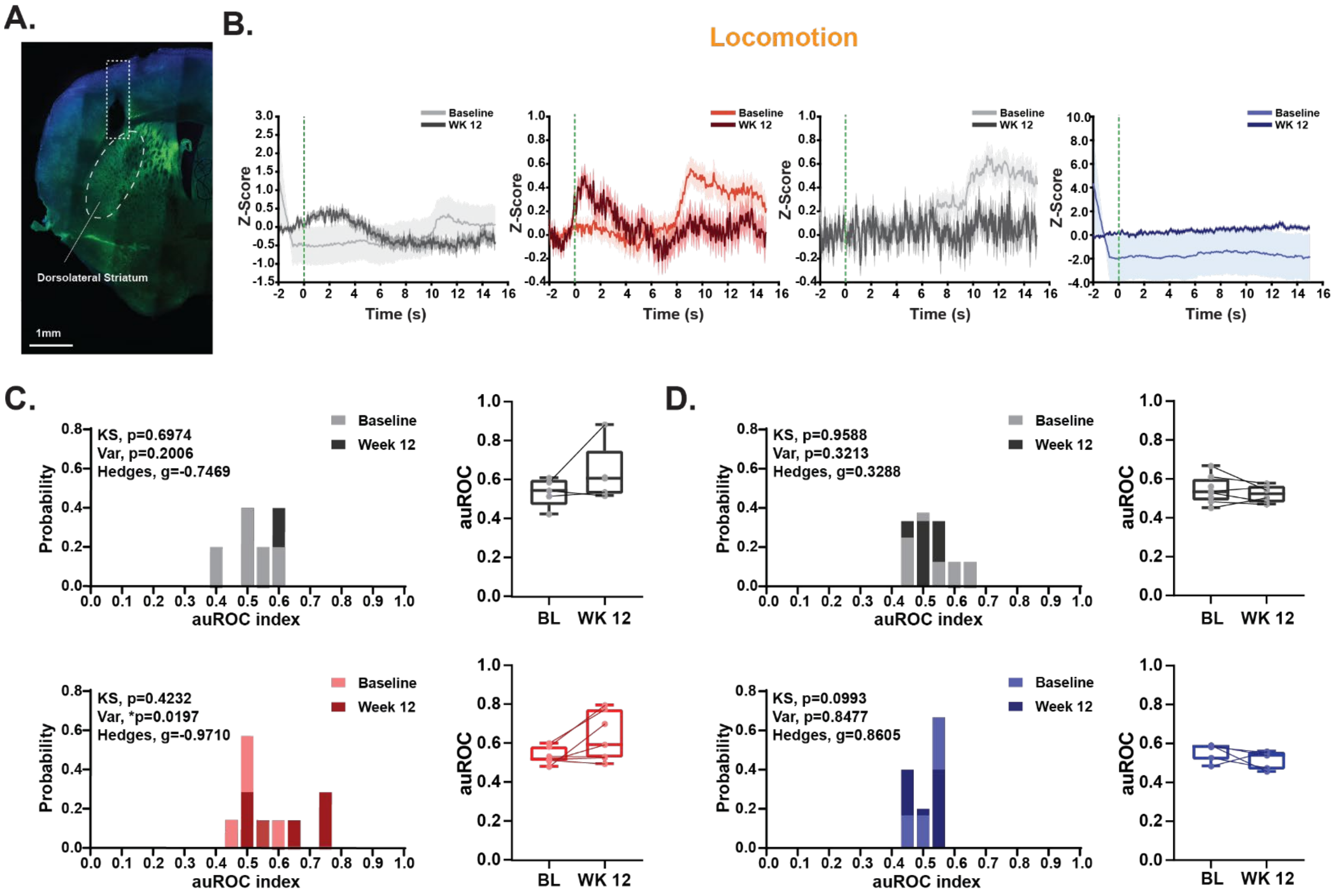

**Supplementary Figure 3**

**(A)** Representative histological image of a fiber photometry mouse. Image shows AAV-GCaMP8s expression (green) with DAPI (blue) in the dorsal striatum with cannula insertion. Scale bar is 1mm.

**(B)** Peri-stimulus time histogram (PSTH) of mean traces sham (gray) and SNI (D1- red, D2- blue) animals comparing baseline (lighter color) to Week 12 (darker color) during locomotive behavior in the Blackbox apparatus. Calcium signals are aligned to the onset of locomotion (t=0). A 1–20s duration filter was applied to the PSTH. Z-score was baseline corrected. The shaded region represents SEM.

**(C)** (*Left*) Probability Histograms of auROC values of D1 activity across baseline compared to week 12 for sham (gray) and SNI mice (red) corresponding to locomotive behavior. (*Right*) auROC values of sham (gray) and SNI (red) of D1 calcium activity from baseline to Week 12. There were no differences in auROC distribution in the Sham or SNI condition across time. There was significance in variance, $F_{(6,6)} = 0.1174$, *p = 0.0197 in the SNI condition but not in the sham condition. Sham: effect size, g =-0.75. N(BL) = 5 and N(WK12) = 5. SNI: effect size, g = -0.97. N(BL) = 7 and N(WK12) = 7.

**(D)** (*Left*) Probability Histograms of auROC values of D2 activity across baseline compared to week 12 of sham (gray) and SNI mice (blue) corresponding to locomotive behavior. (*Right*) auROC values of sham (gray) and SNI (blue) of D2 calcium activity from baseline to Week 12. There were no differences in the sham or SNI condition across time. Sham: effect size, g = -0.33; N(BL) = 8 and N(WK12) = 6. SNI: effect size, g = -0.86; N(BL) = 6 and N(WK12) = 5.

Supplementary Figure 4

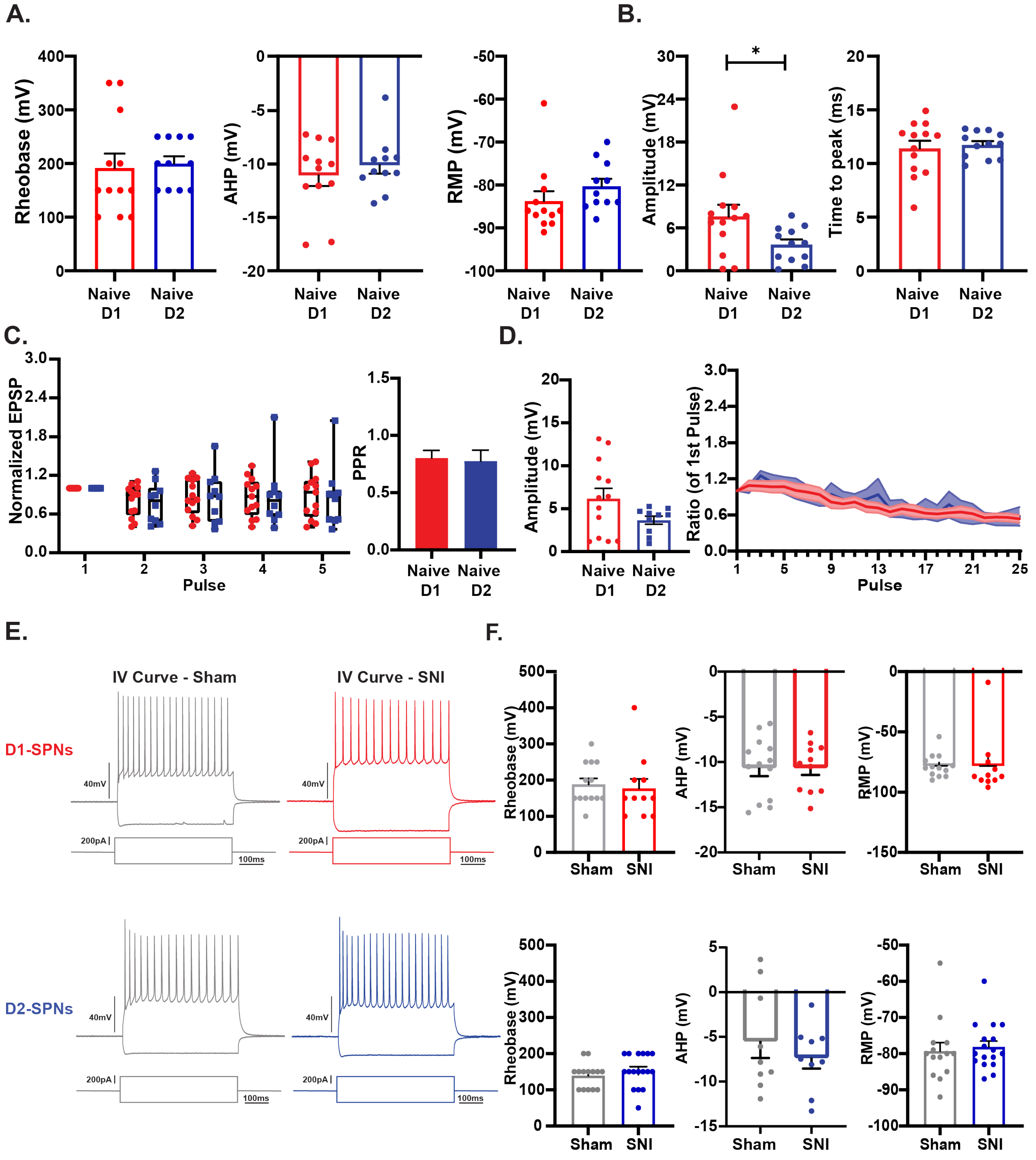

**Supplementary Figure 4**

**(A)** Intrinsic parameters in the naïve condition. There were no significant differences in rheobase, after-hyperpolarization (AHP), and resting membrane potential (RMP) of D1SPNs (red) and D2-SPNs (blue). Unpaired t-test. N = 3 animals for D1 and D2. Error bars represent ± SEM.

**(B)** Quantification of single pulse protocol parameters in naïve condition. (Left) There was a significant difference in PSP amplitude, $t_{(23)} = 2.136$, *p = 0.0435. (*Right*) There were no differences in time to peak of D1-SPNs (red) and D2-SPNs (blue). Unpaired t-test. N = 3 animals for D1 and D2. Error bars represent ± SEM.

**(C)** Quantification of paired-pulse protocol parameters in naïve condition. There were no significant differences in normalized EPSP (*left*) or PPR (right). N = 3 animals for D1 and D2 animals. Error bars represent ± SEM.

**(D)** Quantification of train stimulation protocol parameters in the naïve condition. (*Left*) There were no significant differences in average PSP amplitude. Unpaired t-test. (Right) There were no significant differences in ratio of the 1st pulse. Repeated Two-Way ANOVA with Sidak's multiple comparisons for ratio. Error bars represent ± SEM. N = 3 animals for D1 and D2 animals.

**(E)** Representative current-voltage curves of (*top*) D1-SPNs from sham (gray) and SNI (red) conditions and (bottom) D2-SPNs of sham (gray) and SNI (blue) conditions, during the acute phase.

**(F)** There were no significant differences in intrinsic parameters of rheobase, AHP, and RMP of D1SPNs (red) and D2-SPNs (blue) of both sham and SNI conditions during the acute phase of pain. Two-tail Unpaired t-test. Error bars represent ± SEM. N = 3-4 animals for sham and SNI.

### Supplementary Figure 5

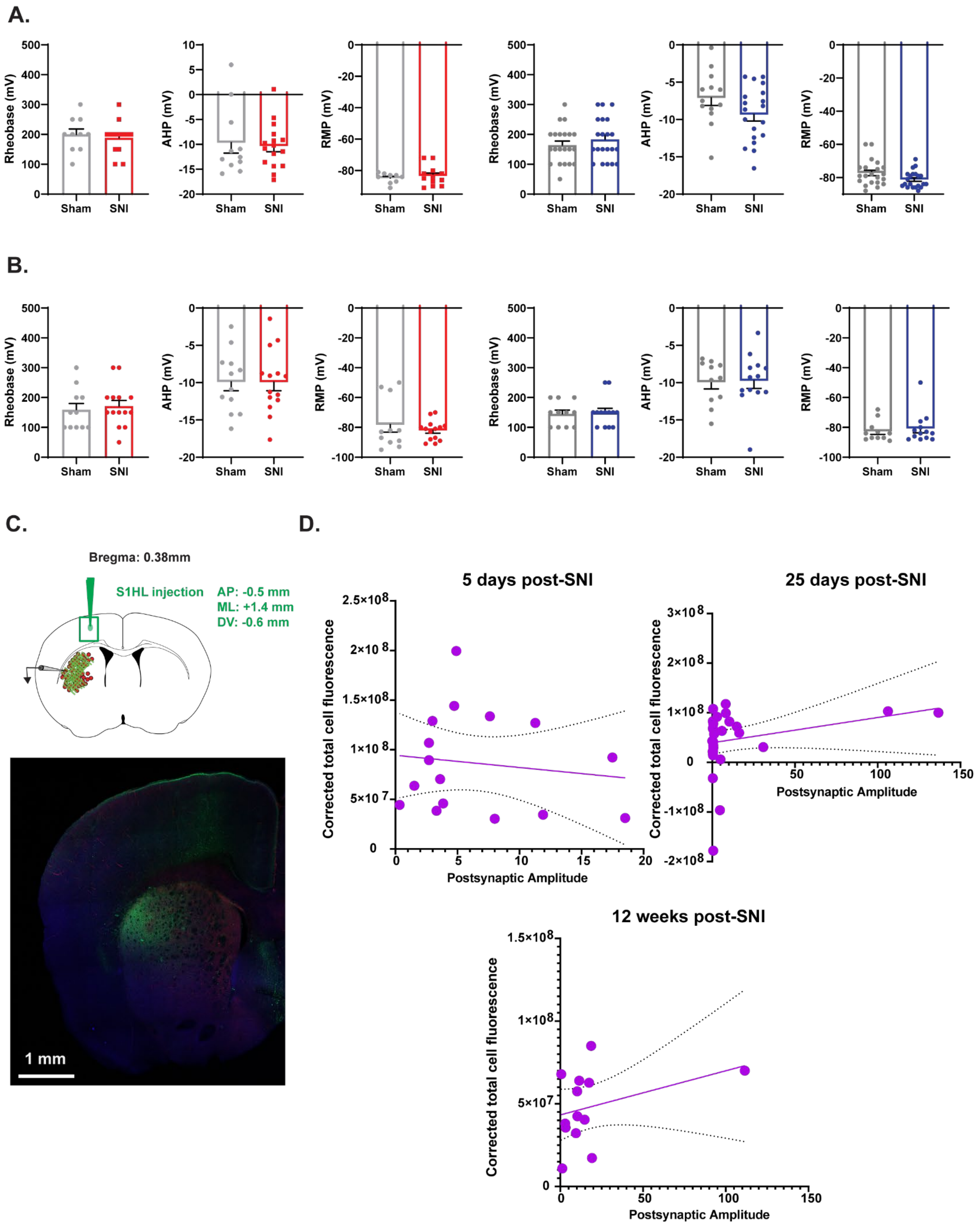

**Supplementary Figure 5**

**(A)** There were no significant differences in intrinsic parameters of rheobase, AHP, and RMP of D1SPNs (red) and D2-SPNs (blue) of both sham and SNI conditions during the transition phase (Week 3) of pain. Unpaired t-test. Error bars represent ± SEM. N = 5-6 animals for sham and SNI.

**(B)** There were no significant differences in intrinsic parameters of rheobase, AHP, and RMP of D1SPNs (red) and D2-SPNs (blue) of both sham and SNI conditions during the late chronic phase of pain (Week 12). Unpaired t-test. Error bars represent ± SEM. N = 3-4 animals for sham and SNI.

**(C)** (*Top*) Schematic EFYP-ChR2 injected into S1HL region Representative histological image of an acute brain slice after postsynaptic recordings from electrophysiological experiments. Image shows ChR2 expression (green) in the dorsal striatum (SPN, Ai14, red) with possible cell fill from postsynaptic recording cell (Biocytin, blue). Scale bar is 1mm.

**(D)** Correlational plot between corrected total cell fluorescence (ChR2) and postsynaptic amplitude for either 5 days-, 25 days-, or 12 weeks-post-injury. There were no significant correlations between viral expression and postsynaptic response. Linear regression. For 5 days post-injury, N = 16; for 25 days post-injury, N = 32; for 12 weeks post-injury, N = 13.

### Supplementary Figure 6

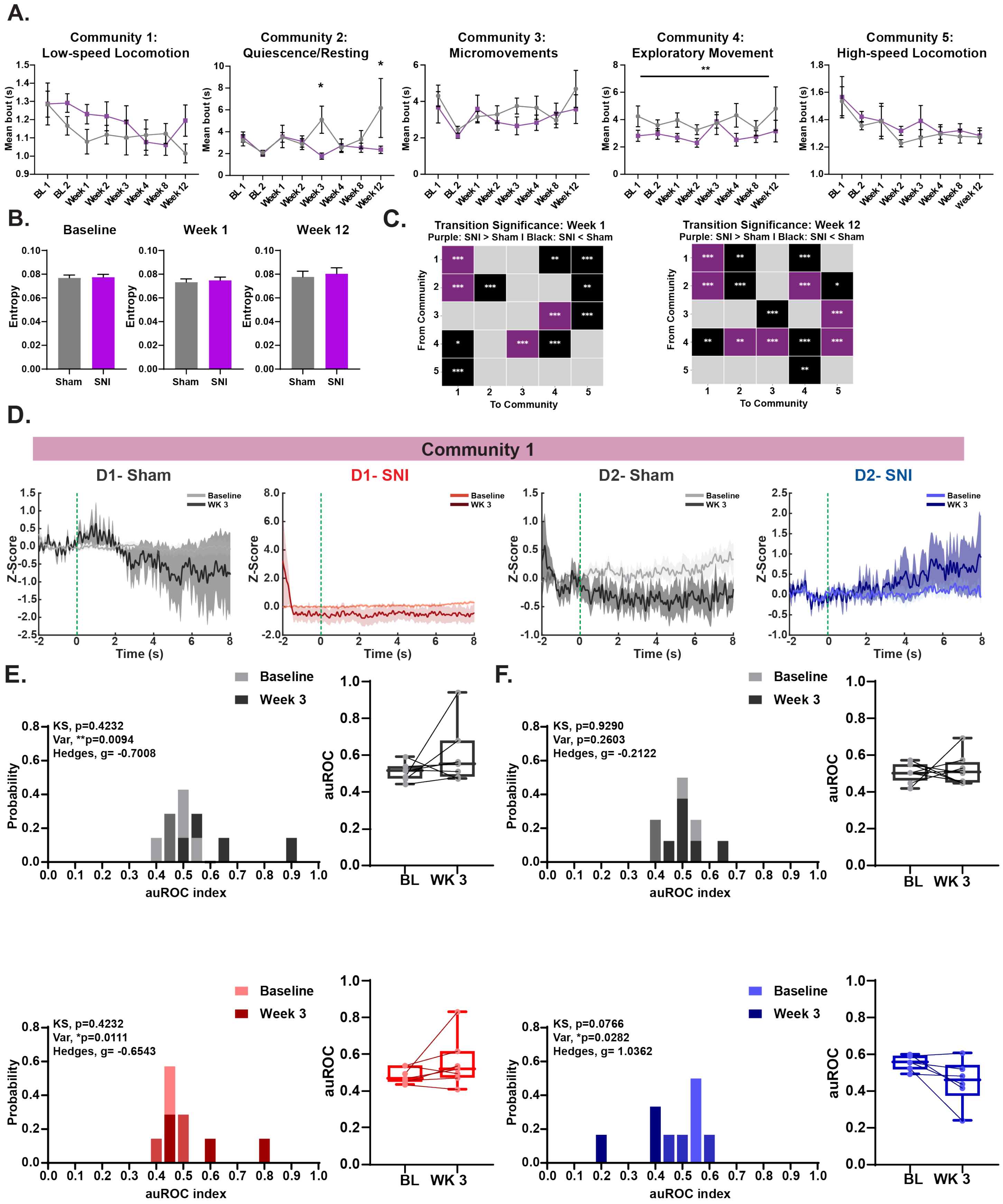

**Supplementary Figure 6**

**(A)** Average bouts of time spent in each community. There were no significant differences in community 1. In community 2, there was a significant interaction $F_{ConditionxTime(1, 202)}$ = 2.066, *p = 0.0488 and significant main effect of condition $F_{condition(1, 202)}$ = 4.970, *p = 0.0269, with significant post-hoc comparisons at week 3 (*p = 0.0278) and week 12 (*p = 0.0192). There were no significant differences in community 1. In community 4, there was a significant main effect of condition $F_{condition(1, 202)}$ = 10.12, **p = 0.0017 without any post-hoc differences. There were no significant differences in community 5. Mixed-effects analysis (REML) with Sidak's multiple comparisons. N = 11-15 for sham and N = 12-13 for SNI.

**(B)** Entropy during baseline, week 1, and week 12. No significant differences were found in any of these timepoints. For baseline, N = 15 for sham and N = 13 for SNI; for week 1, N = 15 for sham and N = 13 for SNI; for week 12, N = 11 for sham and N = 12 for SNI.

**(C)** Reorganized community transition architecture at Week 1 and Week 12. Heatmap depicting pairwise transition significance between the five behavioral communities (C1: Low-speed Locomotion; C2: Quiescence; C3: Micromovements; C4: Exploratory Movement; C5: High-speed Locomotion) at Week 3 post-injury. Rows indicate the originating community ("From"), and columns indicate the terminating community ("To"). Colors indicate the direction of significant probability alterations in SNI relative to Sham: purple indicates significant increase in transition frequency (SNI>Sham), black indicates a significant reduction (SNI<Sham), and gray denotes non-significant transitions (NS). Significance was determined via two-tailed Pearson's $X^2$ contingency tests with Laplace (+1) continuity smooth (*p<0.05, **p<0.01, ***p<0.001).

**(D)** Peri-stimulus time histogram (PSTH) of mean traces of sham (gray) and SNI (D1- red, D2- blue) animals comparing baseline (lighter color) to Week 3 (darker color) during community 2. Shaded region represents SEM.

**(E)** (*Left*) Probability Histograms of auROC values across baseline compared to week 3 of sham (gray) and SNI mice (red) corresponding to community 1. (*Right*) auROC values of sham (gray) and SNI (red) of D1 calcium activity from baseline to Week 3. There were no differences in auROC distribution for both sham and SNI. There were significant differences in variance in Sham, $F_{(7,7)}$ = 0.0882, **p = 0.0094, and in SNI, $F_{(7,7)}$ = 0.0940, *p = 0.0111. Sham: effect size, g = -0.28, N(BL) = 7, N(WK3) = 7. SNI: effect size, g =-0.17, N(BL) = 7 and N(WK3) = 6.

**(F)** (*Left*) Probability Histograms of auROC values across baseline compared to week 3 of sham (gray) and SNI mice (blue) corresponding to community 1. (*Right*) auROC values of sham (gray) and SNI (red) of D2 calcium activity from baseline to Week 3. There were no differences in auROC distribution in either sham or SNI. For SNI, there was a significant difference in variance, $F_{(5,5)}$ = 0.1067, *p = 0.0282. Sham: effect size, g = 0.3965, N(BL) = 8, N(WK3) = 3. SNI: effect size, g =0.0647, N(BL) = 6 and N(WK3) = 5.

### Supplementary Figure 7

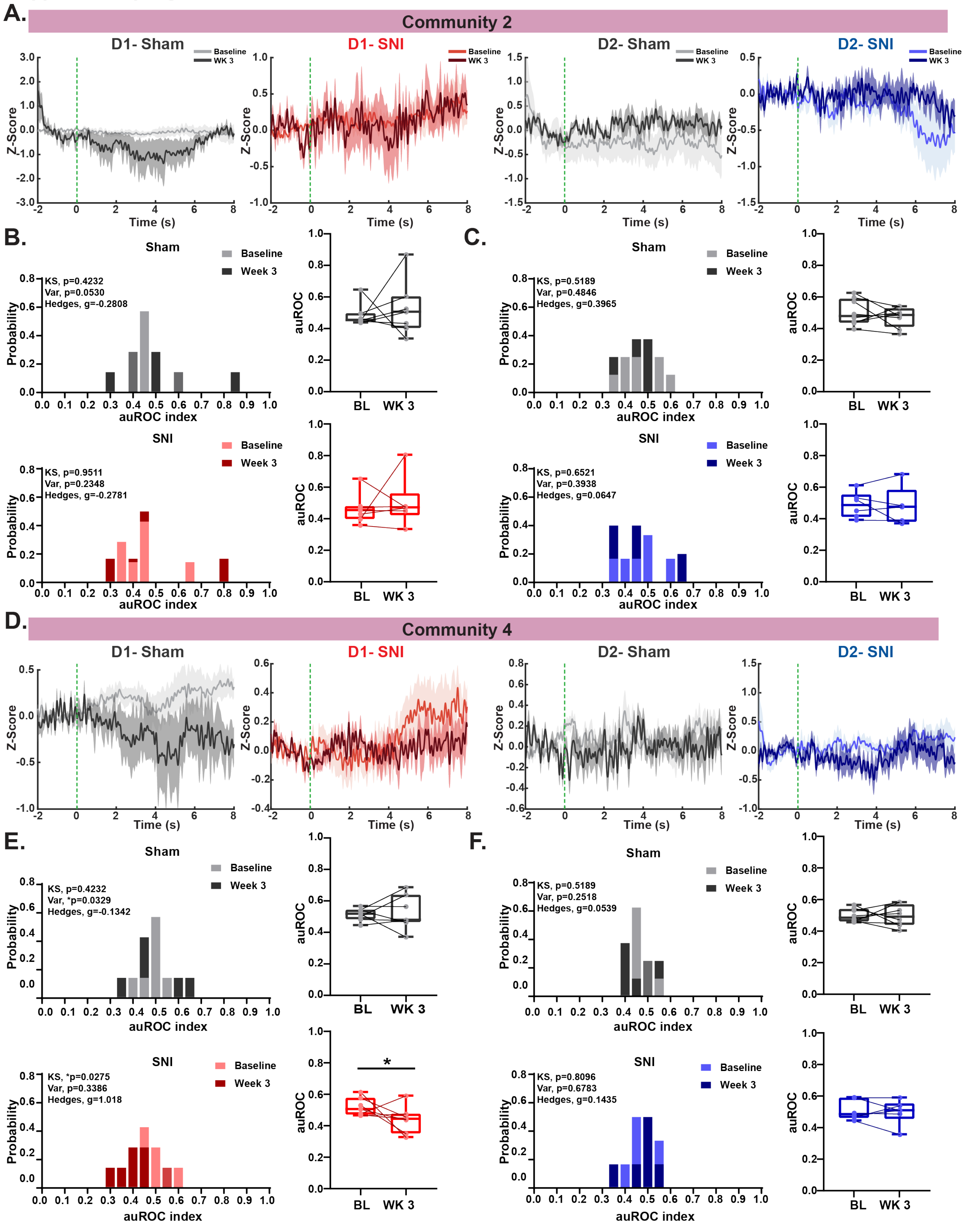

**Supplementary Figure 7**

**(A)** Peri-stimulus time histogram (PSTH) of mean traces sham (gray) and SNI (D1- red, D2- blue) animals comparing baseline (lighter color) to Week 3 (darker color) during community 2. Shaded region represents SEM.

**(B)** (*Left*) Probability Histograms of auROC values across baseline compared to week 3 of sham (gray) and SNI mice (red) corresponding to community 2. (*Right*) auROC values of sham (gray) and SNI (red) of D1 calcium activity from baseline to Week 3. There were no differences in auROC distribution in either sham or SNI. Sham: effect size, g = -0.28, N(BL) = 7, N(WK3) = 7. SNI: effect size, g =-0.17, N(BL) = 7 and N(WK3) = 6.

**(C)** (*Left*) Probability Histograms of auROC values across baseline compared to week 3 of sham (gray) and SNI mice (blue) corresponding to community 2. (*Right*) auROC values of sham (gray) and SNI (red) of D2 calcium activity from baseline to Week 3. There were no differences in auROC distribution in either sham or SNI. Sham: effect size, g = 0.3965, N(BL) = 8, N(WK3) = 3. SNI: effect size, g =0.0647, N(BL) = 6 and N(WK3) = 5.

**(D)** Peri-stimulus time histogram (PSTH) of mean traces sham (gray) and SNI (D1- red, D2- blue) animals comparing baseline (lighter color) to Week 3 (darker color) during community 4. Shaded region represents SEM.

**(E)** (*Left*) Probability Histograms of auROC values across baseline compared to week 3 of sham (gray) and SNI mice (blue) corresponding to community 2. (*Right*) auROC values of sham (gray) and SNI (red) of D1 calcium activity from baseline to Week 3. There were differences in auROC distribution in the SNI condition ($D_{(7,7)}$ = 0.7143, *p =0.0275; KS Test). There were no differences in variance (p = 0.3386). Effect size, Hedge's g = 1.0183. N(BL) = 7, N(WK3) = 7. There were no differences in distribution of auROC values in the sham condition. There was a significant difference in variance, $F_{(6,6)}$ = 0.1443, *p = 0.0329. Effect size, g = -0.1342. N(BL) = 7 and N(WK3) = 7.

**(F)** (*Left*) Probability Histograms of auROC values across baseline compared to week 3 of sham (gray) and SNI mice (blue) corresponding to community 4. (*Right*) auROC values of sham (gray) and SNI (red) of D2 calcium activity from baseline to Week 3. There were no differences in auROC distribution in either sham or SNI. Sham: effect size, g = 0.0539, N(BL) = 8, N(WK3) = 8. SNI: effect size, g =0.1435, N(BL) = 6 and N(WK3) = 6.
